# Comprehensive isotopomer analysis of glutamate and aspartate in small tissue samples

**DOI:** 10.1101/2022.07.31.502208

**Authors:** Feng Cai, Divya Bezwada, Ling Cai, Rohit Mahar, Zheng Wu, Panayotis Pachnis, Chendong Yang, Sherwin Kelekar, Wen Gu, Hieu S. Vu, Thomas P. Mathews, Lauren G. Zacharias, Misty Martin-Sandoval, Duyen Do, K. Celeste Oaxaca, Eunsook S. Jin, Vitaly Margulis, Craig R. Malloy, Matthew E. Merritt, Ralph J. DeBerardinis

## Abstract

Stable isotopes are powerful tools to assess metabolism. ^13^C labeling is detected using nuclear magnetic resonance spectroscopy (NMRS) or mass spectrometry (MS). MS has excellent sensitivity but generally cannot discriminate among different ^13^C positions (isotopomers), whereas NMRS is less sensitive but reports some isotopomers. Here, we develop an MS method that reports all 16 aspartate and 32 glutamate isotopomers while requiring 1% of the sample used for NMRS. This method discriminates between pathways that result in the same number of ^13^C labels in aspartate and glutamate, providing enhanced specificity over conventional MS. We demonstrate regional metabolic heterogeneity within human tumors, document the impact of fumarate hydratase deficiency in human renal cancers, and investigate the contributions of TCA cycle turnover and CO_2_ recycling to isotope labeling in vivo. This method can accompany NMRS or standard MS to provide outstanding sensitivity in isotope labeling experiments, particularly in vivo.

## INTRODUCTION

Many common human diseases, including diabetes, cardiovascular diseases and cancer, are associated with or caused by perturbations of tissue metabolism (DeBerardinis and Thompson, 2012). A thorough characterization of the metabolic state in diseased tissues can improve our understanding of pathophysiology and provide insights for new therapies. Stable isotope tracers including ^13^C, ^2^H and ^15^N are powerful tools to probe metabolism in living systems (Buescher et al., 2015; Jang et al., 2018). Because these isotopes do not undergo radioactive decay, they are considered safe to administer to patients and animal models, and a wide variety of labeled nutrients are readily available for experimental or clinical use. Nutrients labeled with a stable isotope (e.g. ^13^C-glucose) may be introduced to the study subject via enteral or parenteral routes. Metabolism of the labeled nutrient transmits the isotope to metabolic products. Labeling patterns in these products, obtained through fluid or tissue sampling, can be inspected to infer metabolic activity in the intact system. In humans, stable isotope tracing has been used for many years to assess turnover of glucose, lipids and proteins; rates of whole-body substrate oxidation; and alteration of these processes by disease (Kim et al., 2016; Petersen et al., 2003; Shulman et al., 1990; Sunny et al., 2011; Wolfe and Peters, 1987). Several recent studies have assessed human tumor metabolism by examining isotope transfer from labeled nutrients in the circulation to metabolites extracted from tumor specimens (Fan et al., 2009; Faubert et al., 2017; Ghergurovich et al., 2021; Johnston et al., 2021; Maher et al., 2012; Sellers et al., 2015).

Mass spectrometry (MS) and nuclear magnetic resonance spectroscopy (NMRS) are capable of detecting isotope enrichment in metabolites of interest. MS has excellent sensitivity and readily determines both the total enrichment of a metabolite pool and the contributions of each isotopologue to the pool. For example, in the case of a ^13^C labeling experiment designed to examine a metabolite with *i* carbons, MS reports the fraction of the pool contributed by M+0, M+1, M+2 … M+*i* forms of the metabolite, where M is the mass of the unlabeled metabolite. On the other hand, simple assessment of isotopologues by MS does not report the position of ^13^C within the metabolite. This is a significant limitation, because a metabolite of *i* carbons has *i*+1 isotopologues but 2*^i^* labeled forms (isotopomers) when the position of each ^13^C is considered. Therefore, isotopologue analysis by simple MS is informative but lacks the full complement of labeling information afforded by isotopomer analysis. NMRS is the analytical method of choice for isotopomer analysis because the local magnetic environment around an atomic nucleus translates into a predictable position on a chemical shift spectrum, which allows the precise position of ^13^C within a molecule to be reported (Jeffrey et al., 1991). However, NMRS generally does not report all possible isotopomers of a metabolite, and it has low sensitivity relative to MS.

Isotopomer analysis has classically been used to assess the tricarboxylic acid (TCA) cycle, a mitochondrial pathway of fuel oxidation (Malloy et al., 1996). The TCA cycle provides reducing equivalents for oxidative phosphorylation and biosynthetic intermediates for lipid, protein, and nucleotide biosynthesis. This pathway is therefore central to both energy formation and the production of macromolecules. Carbon enters the TCA cycle via the acetyl-CoA pool, which largely arises from pyruvate dehydrogenase (PDH) and fatty acid or ketone oxidation; or through anaplerotic pathways supplied by pyruvate carboxylation and oxidation of some amino acids and other fuels. The relative contributions of different acetyl-CoA sources, anaplerosis, pyruvate recycling, cycle turnover and other activities to overall flux are encoded by specific positions of ^13^C within TCA cycle intermediates, emphasizing the power of isotopomer analysis in analyzing the TCA cycle. Glutamate isotopomers are particularly useful because glutamate provides information about the acetyl-CoA and oxaloacetate (OAA) that supply citrate synthase, and because the high intracellular abundance of glutamate makes it a convenient metabolite for analysis.

Several previous studies have used fragmentation patterns in tandem mass spectrometry to examine isotopomers of metabolites related to the TCA cycle. Gas chromatography-tandem mass spectrometry (GC-MS/MS) was used to partially resolve isotopomers of glutamate extracted from hearts perfused with various ^13^C-labeled fuels and to calculate anaplerotic flux and acetyl-CoA enrichment (Jeffrey et al., 2002). GC-MS/MS was also used to derive all 16 aspartate isotopomers, which were then validated against a series of ^13^C-labeled aspartate standards (Choi et al., 2012). Grouped rather than individual isotopomers from multiple TCA cycle intermediates, including some groups of glutamate isotopomers, were determined with liquid chromatography-tandem mass spectrometry (LC-MS/MS) and used to calculate fluxes related to central carbon metabolism in cultured cells (Alves et al., 2015).

We set out to develop an LC-MS/MS method to identify as many of the 32 individual isotopomers of glutamate and 16 individual isotopomers of aspartate as possible. Our motivation to develop this technique is the increasing use of stable isotope tracers to assess cancer metabolism *in vivo*. These studies produce complex labeling patterns that are difficult to interpret without positional assignment of ^13^C. The large sample size required for NMRS makes it impractical to use this technique to study regional heterogeneity in ^13^C labeling, which is significant in human solid tumors. We benchmarked the new method against NMRS and find that it provides similar information about isotopomer distributions despite requiring only about 1% of the sample. We also demonstrate the new method’s superiority to simple isotopologue analysis in assessing metabolism of cultured cancer cells and tumors growing in mice and patients.

## RESULTS

### Development and validation of LC/MS isotopomer analysis approaches

We first developed an approach to detect all 32 (i.e. 2^5^) isotopomers of glutamate using multiple reaction monitoring (MRM) with LC-MS/MS. Glutamate contains five carbons and four carbon-carbon σ bonds. We examined the fragmentation pattern of glutamate and found seven precursor/product ion pairs (M/m) as summarized in Fig.1A, in which precursor ions are denoted with a capital M and product ions are denoted with a lowercase m. Two ion pairs, 146/41 and 146/59, report the C4-5 fragment, but the 146/41 ion pair produces a higher-quality peak on a Sciex QTRAP 6500. We use the 146/41 ion pair in the analyses described below. The 146/74 and 148/84 (or 148/102) ions report the C1-2 and C2-5 fragments, respectively. Both 148/84 and 148/102 positive ions resulted from C1-C2 cleavage and showed high quality peaks, but 148/84 was chosen because it provided more consistent signals. The 146/102 negative ion is a mixture of C1-4 and C2-5 fragments, both resulting from loss of a CO_2_ during glutamate fragmentation. By examining the 148/104 and 148/103 ions fragmented from a [1,2-^13^C]glutamate standard (denoted as glutamate-11000 throughout this paper, with 1 indicating a ^13^C carbon and 0 indicating a ^12^C carbon), we determined that the C1-4 and C2-5 fragments are generated at a fixed ratio of 19:1, allowing relative quantitation of these two fragments (Fig. 1C, E). (Kappelmann et al., 2017). The 148/56 ion mainly arises from the C2-4 fragment along with a small amount (2%) of the C3-5 fragment, which was considered negligible. These fragments were validated by examining the 150/57 and 150/56 fragmentation of a glutamate-11000 standard (Supplementary Data I, Scheme 1a).

**Figure 1.**
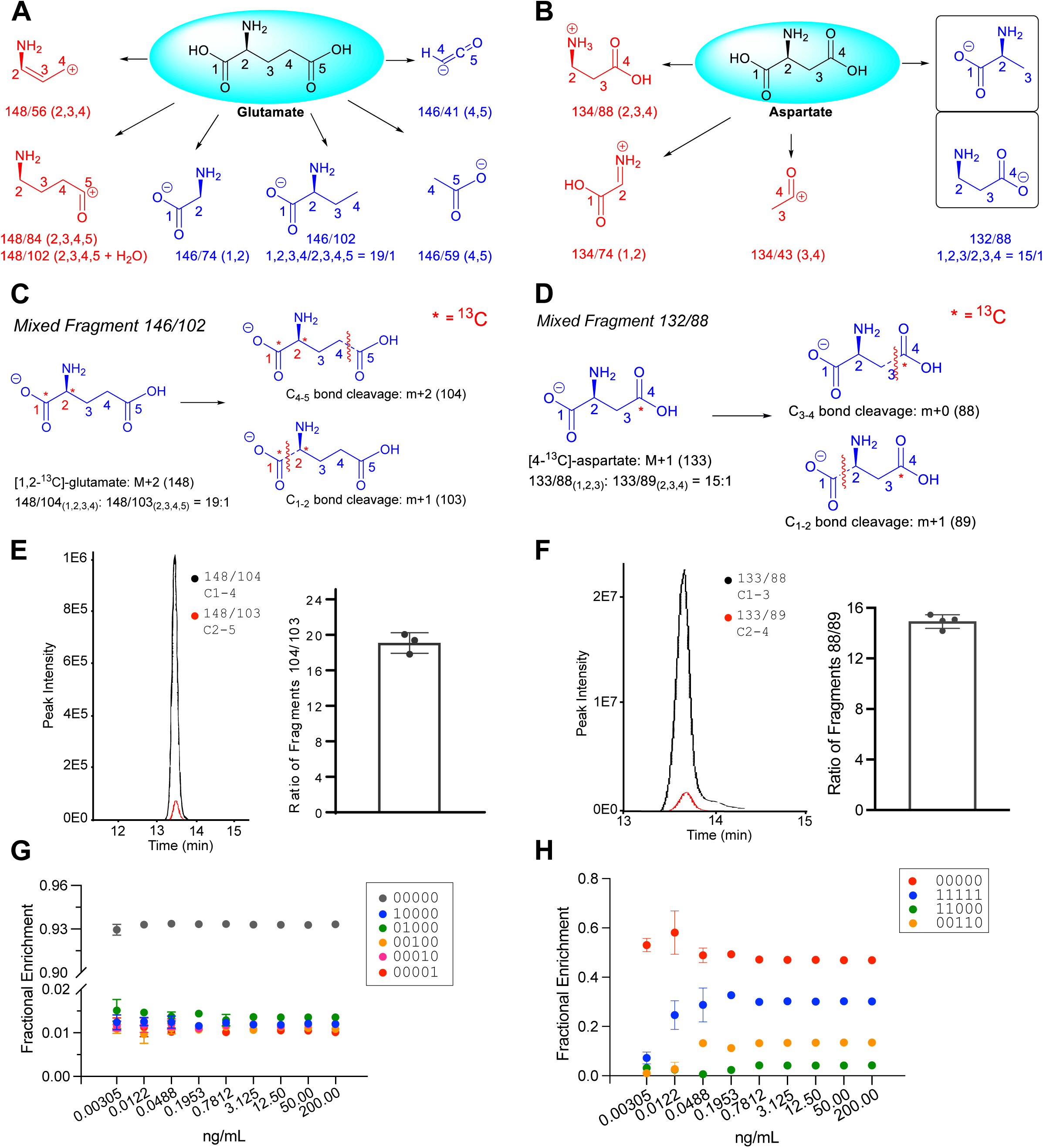
Development of LC-MS/MS analysis of glutamate and aspartate isotopomers. (A-B) LC-MS/MS fragmentation of glutamate (A) and aspartate (B). Carbon numbers correspond to positions in the unfragmented molecule. Parent/daughter ion pairs are described in the text. (C) Composition of the mixed 146/102 fragment from glutamate (D) Composition of the mixed 132/88 fragment from aspartate. (E) Relative abundance of the 148/104 (C1-4) and 148/103 (C2-5) ion pairs from a [1,2-^13^C]glutamate standard (F) Relative abundance of the 133/88 (C1-3) and 133/89 (C2-4) ion pairs from a [4-^13^C]aspartate standard. (G) Relative quantitation of naturally-occurring glutamate isotopomers. (H) Relative quantitation of isotopomers from a mixture of [1,2-^13^C]glutamate (5%); [3,4-^13^C]glutamate (15%); [U-^13^C]glutamate (30%); and unlabeled glutamate (50%), across a range of concentrations.

It is worth noting that the 146/41 ion pair can also be generated from the C2-4 fragment (C_3_H_5_^-^) and may interfere with the C4-5 fragment (C_2_HO^-^) (Kappelmann *et al*., 2017). The contribution of C2-4 versus C4-5 in the 146/41 ion pair can be distinguished by measuring the 151/43 (M+5/m+2, ^13^C_2_HO^-^, C4-5 fragment) and 151/44 (M+5/m+3, ^13^C_3_H_5_^-^, C2-4 fragment) ion pairs from a glutamate-11111 standard (Supplementary Data I, Scheme 1b). We observed both 151/43 and 151/44 on a Sciex QTRAP 5500, but the 151/44 ion pair was below the detection limit on a Sciex QTRAP 6500. If both ion pairs are detected, the ratio of the two ion pairs should be considered in the isotopomer distribution matrix. We also used a parallel reaction monitoring approach on an Orbitrap detector to assess these specific ion pairs, although the remaining data reported in this paper were generated using the MRM method.

With these five main precursor/product ion pairs (146/41, 146/74, 146/102, 148/56, and 148/84) in unlabeled glutamate, additional ion pairs can differentiate labeled higher order isotopomers in the format M+*i*/m+*j* (*j*≤*i*), in which *i* denotes the number of ^13^C in the precursor ion and *j* denotes the number of ^13^C in the product ion. Taking all the ^13^C isotopomers into account, 88 total ion pairs were identified. There are twenty ion pairs associated with labeled forms of 146/102, which mainly represent the C1-4 fragment, and 148/56, which includes the C2-4 fragment and therefore also reports information about C4-C5 bond cleavage. These twenty ion pairs seem redundant, but this complementary information can improve accuracy in some samples. For example, the chromatogram peak for 147/75, an M+1/m+1 labeled form of 146/74, often overlaps with another metabolite. Although this prevents precise analysis of the 147/75 ion pair, other ion pairs can provide complementary information (Supplementary Data I).

These five ion pairs provide information about all four glutamate C-C bond cleavage sites. However, the distribution matrix that linearly maps the 88 ion pairs to all 32 glutamate isotopomers only has a rank of 28 out of a full rank of 32. Obtaining additional fragments of C2-3 and C3-4 may produce a complete rank, but this is not feasible due to the chemical properties of glutamate. Gratifyingly, nonnegative least square regression helps in this situation to obtain information about all 32 glutamate isotopomers. To assess the accuracy of the nonnegative least square regression, a simulation study is provided in Supplemental Data I with the median and 95% confidence interval for isotopomer distribution error based on the simulation. Eighteen of the glutamate isotopomers are highly precise with absolute errors less than 7e-17. The remaining 14 glutamate isotopomers have absolute errors up to +/-3%. However, the 14 isotopomers with the highest apparent uncertainty are all in the groups of M+2 or M+3 isotopologues, and 10 of these 14 contain only single labels in their C4-5 fragments. These isotopomers are relatively scarce in tracing experiments that produce [1,2-^13^C]acetyl-CoA (e.g. [U-^13^C]glucose and [1,2-^13^C]acetate). Two additional isotopomers (glutamate 10011 and 01011) are downstream of OAA 0001,1001, 0010, or 1010. These isotopomers are also expected to be scarce when the major labeled form of acetyl-CoA is [1,2-^13^C]acetyl-CoA (Supplementary Figure 1A). The last two isotopomers, glutamate 10100 and 01100, provide important information related to TCA cycle turnover but have relatively high errors. However, these two isotopomers are directly proportional to glutamate 10111 and 01111 and the fraction of acetyl-CoA supplying the TCA cycle that is labeled as [1,2-^13^C]acetyl-CoA (F_C3_, the fraction of acetyl-CoA ^13^C containing ^13^C at both the carbonyl and methyl carbons). Thus, these last two isotopomers can be evaluated by combining an analysis of glutamate 10111 and 01111 with F_C3_.

We performed a similar isotopomer analysis of aspartate. The fragment ions for aspartate are illustrated in Fig. 1B. We identified 4 ion pairs (134/88, 134/74, 134/43 and 132/88) to distinguish the C-C bond cleavage sites in aspartate. The 132/88 ion pair is a mixture of fragments C1-3 and C2-4 (Fig. 1D). Using an aspartate 0001 standard, we measured the ratio of 133/88 and 133/89 to differentiate the contribution of C1-3 versus C2-4 in ion pair 132/88 and determined that the average ratio of C1-3:C2-4 is 15:1 (Fig. 1F). Similarly, there are twelve ion pairs associated with labeled forms of 134/43 and these ion pairs provide redundant information to improve accuracy in some samples. We repeated similar calculations for all 16 aspartate isotopomers and evaluated their accuracy (Supplementary Data I). Only 4 isotopomers including aspartate-1010, 0110, 1001, and 0101 produced noticeable errors (up to +/-4%). These four isotopomers are not expected to appear at high levels in tracing experiments that produce [1,2-^13^C]acetyl-CoA.

We validated the glutamate isotopomer method using glutamate standards. We first examined a naturally occurring glutamate standard across a range of concentrations without natural abundance correction (Fig. 1G, data in Supplemental Data II). This revealed the presence of all five forms of singly-labeled glutamate (at positions C1-C5) at approximately 1% each, which is close to the expected 1.1% natural abundance of ^13^C at each position. As expected, unlabeled glutamate was approximately 94% (Fig. 1G). We next examined a mixture of 10% glutamate 00000, 15% glutamate 11000 (99% isotope purity, 98% chemical purity), 25% glutamate 00110 (99% isotope purity, 98% chemical purity) and 50% glutamate 11111 (99% isotope purity, 98% chemical purity). Different concentrations of this mixture were used to explore the dynamic range of the method (Fig. 1H). In each sample, all 32 isotopomers were analyzed. When the glutamate concentration exceeds 0.78 ng/mL, the calculated isotopomer fractions match the true fraction. Lower concentrations result in poor signals of some key ion pairs (e.g. 146/41, 146/74) and fail to produce satisfactory results. Isotopomer distributions were accurately reported at concentrations as high as 200 ng/mL, although at this concentration the peak areas exceed the linear curve due to source saturation. As a result, experiments should use at least three concentrations to evaluate whether the concentration is in the linear range. We note that some degree of saturation at the ion source interferes with glutamate ionization but does not affect the ratios among different isotopomers (Liu et al., 2011; Wei et al, 2020). However, isotopomer ratios are altered when the mass spectrometer detector becomes saturated, so the concentration of the highest single isotopomer should be carefully monitored. Practically speaking, these detection limits translate to cell culture samples from a minimum of a 60-mm diameter up to a 150-mm diameter cell culture dish at 80% confluence (roughly 1.5 million – 10 million cells), or roughly 10 mg pieces of human or mouse tumor tissue.

### Validating LC-MS/MS method against conventional NMRS isotopomer analysis

To extend the isotopomer method to biological samples and benchmark it against ^13^C NMR, we cultured H460 lung cancer cells in medium containing [U-^13^C]glucose for 24 h and then extracted the metabolites from three 150-mm diameter dishes at approximately 90% confluence. We used 1% of the extract for LC-MS/MS, and the rest was used for NMRS. ^13^C NMR reported the relative abundance of a subset of glutamate and aspartate isotopomers based on spin coupling generated by adjacent ^13^C nuclei (Fig. 2A,B), whereas LC-MS/MS analysis reported all 32 isotopomers of glutamate and all 16 isotopomers of aspartate (Fig. 2C,D; data in Supplementary Data II). To directly compare the isotopomer distributions reported by the two methods, data from LC-MS/MS were left uncorrected for naturally occurring isotopes and were summarized in the same format as the ^13^C NMR data (Fig. 2E; data in Supplementary Data II). This revealed excellent consistency for isotopomers related to carbons C2-4, which encode most of the information relevant to TCA cycle turnover in this labeling scheme. Somewhat larger discrepancies were observed with C1. These discrepancies could likely be reduced with more sensitive mass spectrometers. We performed a similar comparison of aspartate isotopomers in the same sample (Fig. 2B, D and F). Again, the isotopomer distributions were similar between the two methods. C1-C3 labeling was nearly identical, with somewhat larger discrepancies related to C4 (Fig. 2F).

**Figure 2.**
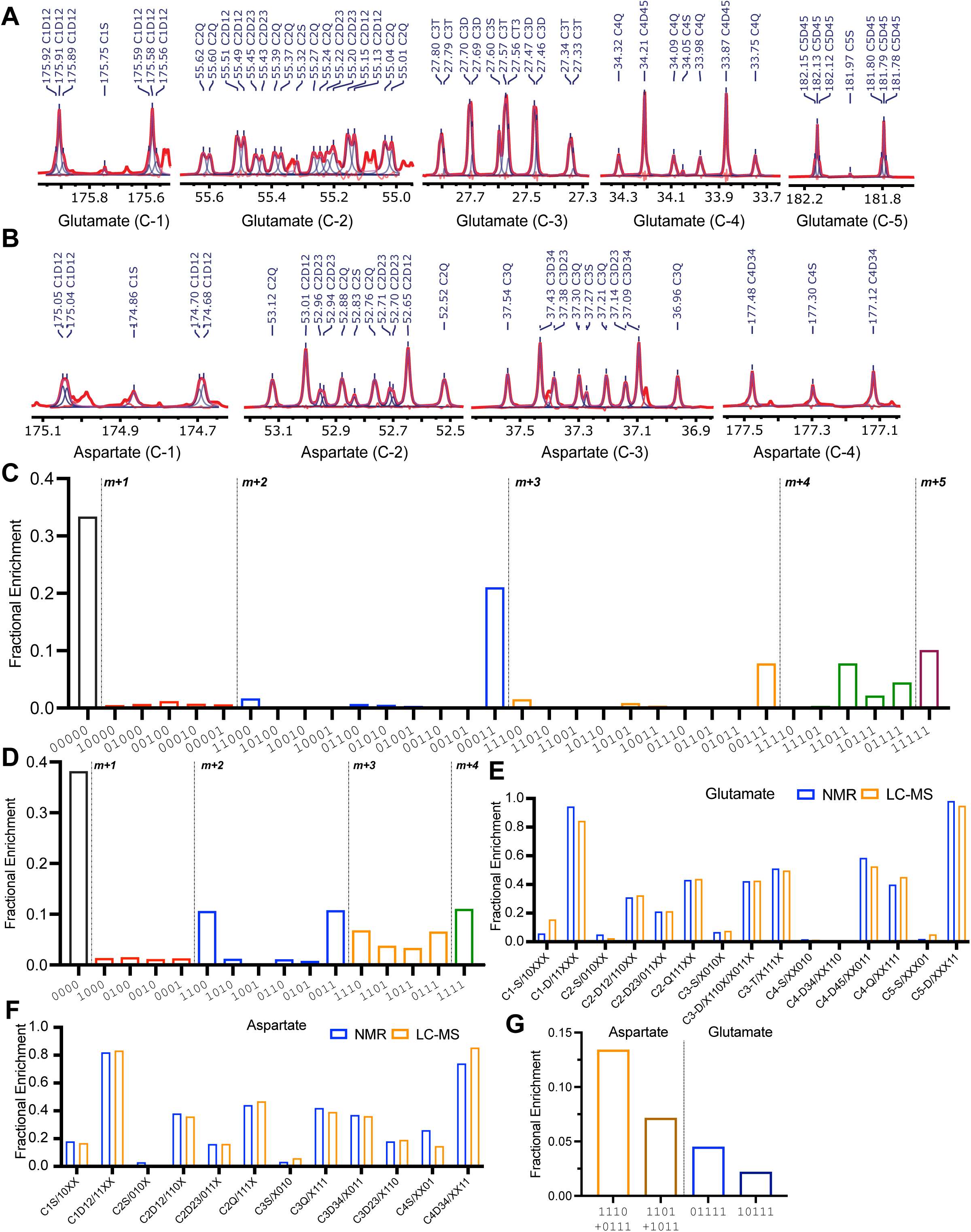
Comparison of isotopomer analysis from NMRS and LC-MS/MS. (A) 1D ^13^C NMR spectrum from H460 lung cancer cells cultured with [U-^13^C]glucose. Multiplets from all five glutamate carbons are displayed. (B) 1D ^13^C NMR spectrum from the same culture as in (A), displaying multiplets from all four aspartate carbons. (C) Relative abundance of 32 glutamate isotopomers by LC-MS/MS. (D) Relative abundance of 16 aspartate isotopomers by LC-MS/MS (E) Comparison of glutamate isotopomer analysis by ^13^C NMR and LC-MS/MS, using data from (A) and (C). (F) Comparison of aspartate isotopomer analysis by ^13^C NMR and LC-MS/MS, using data from (B) and (D).

An advantage of isotopomer analysis is that it allows the calculation of metabolic parameters such as F_C3_ and anaplerosis (y). Using the samples from this [U-^13^C]glucose culture, we compared F_C3_ and y values derived from NMRS and MS. From the LC-MS/MS data, F_C3_ was calculated with an established formula commonly used in NMRS (eq. 1) or by directly using two isotopomers, glutamate 11000 and 11011 (eq. 2). These equations calculated F_C3_ as 0.823 and 0.852, respectively. Eq. 1 also calculated F_C3_ as 0.845 from the NMRS data. Using an equation developed for NMRS (eq. 3), relative anaplerotic flux (y) was calculated as 0.47 from LC-MS/MS data and 0.52 from NMRS data (data in Supplementary Data II). Therefore, the new LC-MS/MS analysis provides similar information about metabolism as the established NMRS approach while requiring only 1% of the input material.

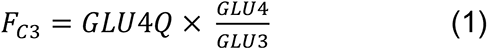

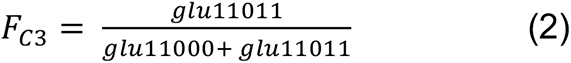

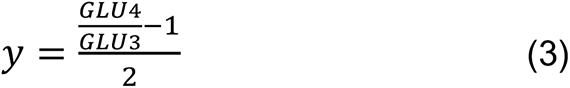

It is significant that the fractions of aspartate 1110 and 0111 differ from aspartate 1101 and 1011 (Fig. 2D). In a simple metabolic network with Ac-CoA as the only carbon source for the TCA cycle, these four m+3 isotopologues should be present in equal fractions because of the symmetric structure of succinate and fumarate (Supplementary Figure 1A). However, ^13^C enters the TCA cycle as both Acetyl-CoA and OAA in cells with concomitant pyruvate dehydrogenase and carboxylase activity (PDH and PC, Supplementary Figure 1B). The difference between the sum of aspartate 1110 and 0111, which arise initially from PC, and aspartate 1101 plus 1011, which arise from multiple pathways, indicates that PC is active and contributes to the TCA cycle (Fig. 2G). The corresponding difference is found in glutamate, in which glutamate 01111 (arising from oxaloacetate 1110 and [1,2-^13^C]acetyl-CoA) is more abundant than glutamate 10111 (arising from oxaloacetate 1101 and [1,2-^13^C]acetyl-CoA). Together these findings emphasize that positional ^13^C assignment by LC-MS/MS detects the consequences of multiple routes of ^13^C entry into the TCA cycle, even among isotopomers with the same number of ^13^C nuclei.

### Detection of distinct modes of anaplerosis in cancer cell lines and mouse tissues

Cancer cells in culture have substantial anaplerotic fluxes as described above for H460 cells. These fluxes allow cells to maintain consistent levels of TCA cycle intermediates while withdrawing intermediates from the cycle to supply biosynthetic pathways. Glutamine catabolism and PC are two well-characterized modes of anaplerosis in cultured cancer cells (Cheng et al., 2011; Yang et al., 2014). We previously analyzed ^13^C enrichment using NMRS and non-positional MS to characterize anaplerosis in SFxL glioma (high glutamine catabolism) and Huh7 hepatoma (high PC) cells (Yang et al., 2014). To test whether the LC-MS/MS method detects these differences, we cultured both cell lines with [U-^13^C]glucose and analyzed positional labeling with natural abundance correction. Over a time course of several hours, Huh7 cells displayed enhanced fractional accumulation of PC-dependent isotopomers of aspartate (1110 and 0111) and glutamate (11100 and 01100) (Fig. 3A-D). Huh7 cells also rapidly accumulated higher-ordered labeling in glutamate (01111 and 11111), in which the former arose from OAA 1110 and [1,2-^13^C]acetyl-CoA, and the latter from OAA 0111 and [1,2-^13^C]acetyl-CoA (Fig. 3B, D).

**Figure 3.**
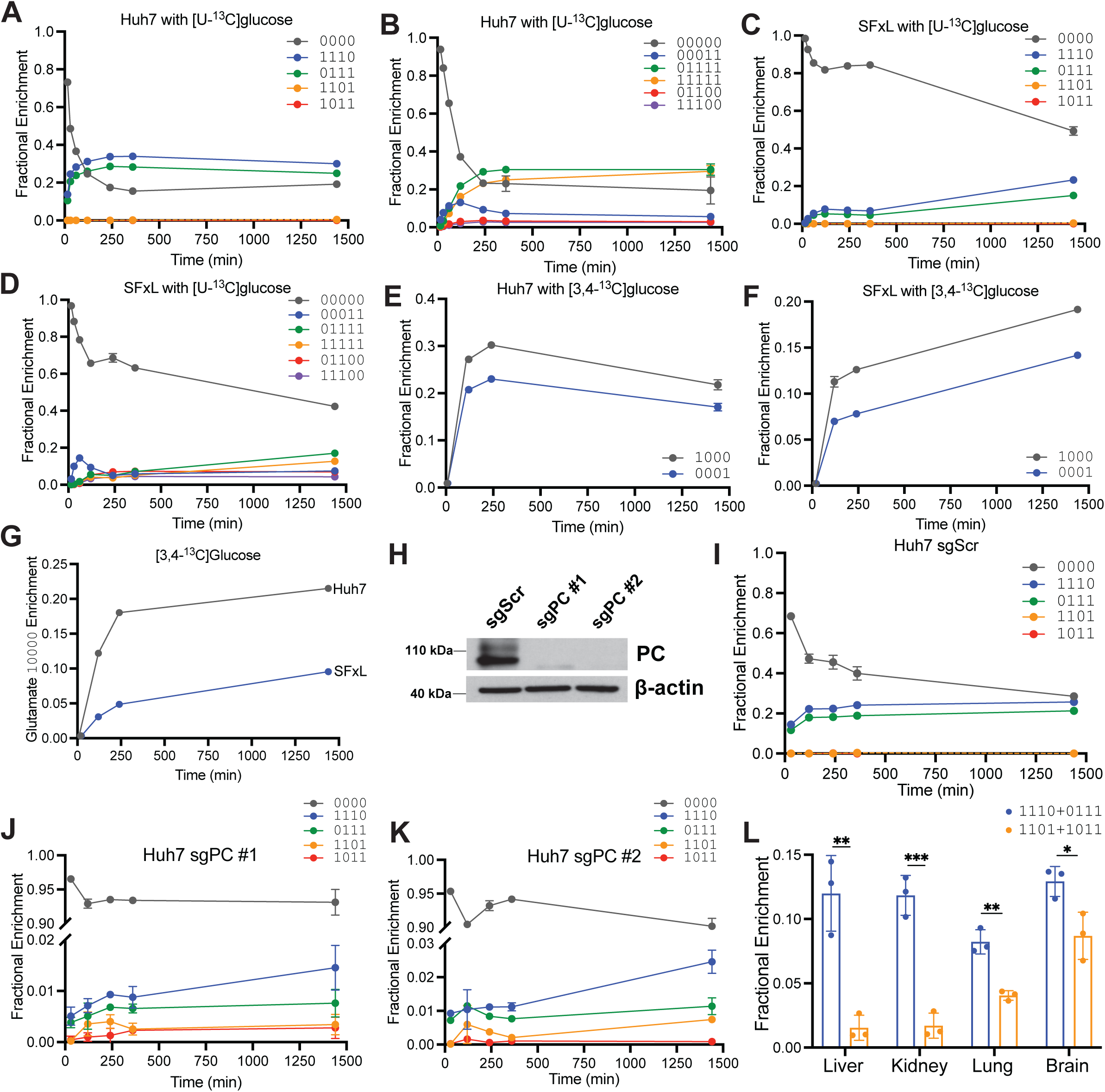
Kinetic analysis of glutamate isotopomers in cancer cells. (A-B) Time course of selected aspartate and glutamate isotopomers in Huh7 hepatoma cells cultured in medium containing [U-^13^C]glucose. (C-D) Time course of selected aspartate glutamate isotopomers in SF-xL gliomas cells cultured in medium containing [U-^13^C]glucose. (E-G) Time course of selected aspartate and glutamate isotopomers in Huh7 hepatoma cells and SF-xL gliomas cells cultured in medium containing [3,4-^13^C]glucose. In panels A-G, enrichment in glutamate and aspartate was assumed to be 0.0 at time 0. (H) Western blot of PC abundance in Huh7 hepatoma cells expressing control guide RNAs (sgScr) or guide RNAs targeting PC (sgPC #1, #2). (I-K) Time course of selected aspartate isotopomers in sgScr and sgPC Huh7 hepatoma cells cultured in medium containing [U-^13^C]glucose. (L) Selected aspartate M+3 isotopomers in mouse organs after infusion with [U-^13^C]glucose. P values: ns = P > 0.05; * =P ≤ 0.05; ** = P ≤ 0.01; *** = P ≤ 0.001; **** = P ≤ 0.0001. Unpaired t tests were used and the data are shown as mean ± standard deviation

To further examine PC-dependent labeling, we used [3,4-^13^C]glucose. With this tracer, ^13^C enters the TCA cycle only via OAA as a consequence of PC activity, but not via acetyl-CoA (Supplementary Fig. 2). In Huh7 cells, labeled aspartate appeared rapidly as 1000 and 0001, whereas these i11sotopomers accumulated slowly in SFxL cells (Fig. 3E,F). Glutamate 10000, which also arises downstream of PC, also appeared more rapidly in Huh7 cells (Fig. 3G).

We next examined the effects of PC loss in Huh7 cells by depleting PC using CRISPR-Cas9 (Fig. 3H). During culture with [U-^13^C]glucose, control cells expressing a non-targeting guide RNA (sgScr) demonstrated rapid production of aspartate 1110 and 0111, similar to parental Huh7 cells (Fig. 3I). However, in the two PC-deficient lines, labeled forms of aspartate were reduced 10-fold or more (Fig. 3J,K).

Finally, we explored PC activity in several mouse organs after infusing mice with [U-^13^C]glucose, extracting metabolites and examining M+3 labeling in aspartate. In the liver and kidney, the sum of aspartate 1110 plus 0111 far exceeded the sum of aspartate 1101 plus 1011, indicating PC activity (Fig. 3L). A smaller but still positive difference was observed in lung and brain. These data are consistent with the known robust PC activity in liver and kidney.

### Application of LC-MS/MS to interpret labeling patterns in a mouse model of tumor growth

We next used LC-MS/MS isotopomer analysis to assess labeling in tumors from mice infused with ^13^C-labeled nutrients. In these experiments, our goals were to provide clarity about the labeling patterns arising in tumor metabolites, and to test whether the LC-MS/MS method detects labeling changes induced by a metabolic inhibitor. We generated xenografts from SK-N-AS neuroblastoma cells in immunodeficient NSG mice, infused the mice with either [U-^13^C]glucose or [1,2-^13^C]acetate, and treated them with a vehicle control (DMSO) or the compound IACS-010759, an inhibitor of Complex I in the electron transport chain (ETC). Because Complex I recycles NADH to NAD+, and PDH requires NAD+ as a cofactor, IACS-010759 reduces the fractional abundance of isotopomers arising from PDH (Pachnis et al, 2022). Indeed, glutamate 00011 was suppressed by IACS-010759 during infusion with [U-^13^C]glucose (Fig. 4A). IACS-010759 reduced F_C3_ from 0.32 to 0.19 in these xenografts (Fig. 4B). Labeling in the second turn of the TCA cycle (e.g. glutamate 11000) was also suppressed by IACS-010759 (Fig. 4C). IACS-010759 increased rather than decreased glutamate 00011 and F_C3_ after infusion with [1,2-^13^C]acetate, which does not require PDH to label Ac-CoA (Fig. 4D,E). Therefore, the method detects the anticipated effects of Complex I inhibition in tumor-bearing mice.

**Figure 4.**
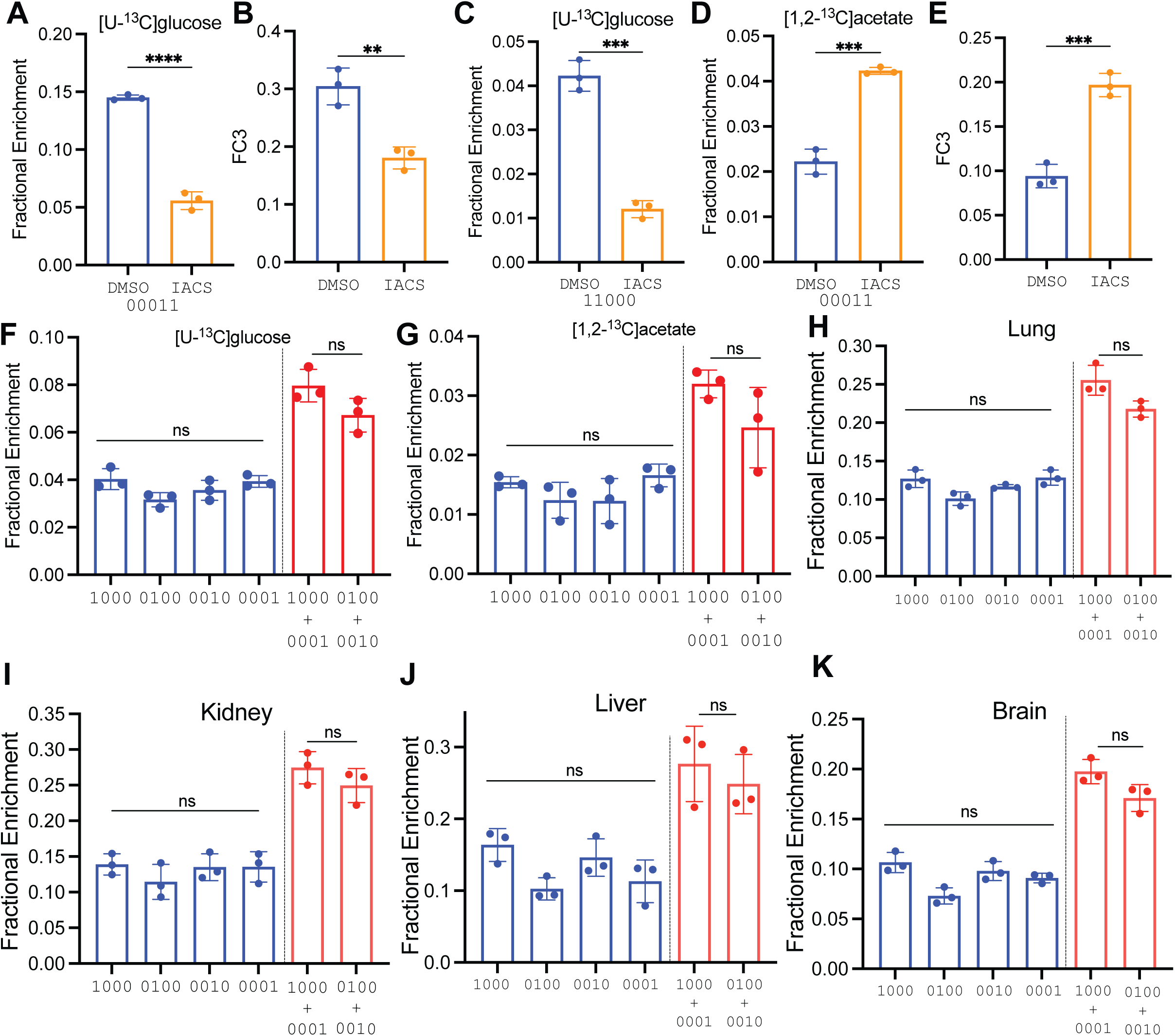
In vivo applications of LC-MS/MS isotopomer analysis. (A-C) Relative [4,5-^13^C]glutamate abundance (A), F_C3_ fraction (B) and abundance of [1,2-^13^C]glutamate as a fraction of the total glutamate pool in neuroblastoma xenografts after infusion with [U-^13^C]glucose and treatment with DMSO or IACS-010759. (D-E) Relative [4,5-^13^C]glutamate abundance (D) and F_C3_ fraction (E) in neuroblastoma xenografts after infusion with [U-^13^C]glucose and treatment with DMSO or IACS-010759. (F-G) Aspartate isotopomers from DMSO-treated neuroblastoma xenografts after infusion with [U-^13^C]glucose (F) or [1,2-^13^C]acetate (G). (H-K) Aspartate M+1 isotopomers in mouse organs after infusion with [U-^13^C]glucose. P values: ns = P > 0.05; * =P ≤ 0.05; ** = P ≤ 0.01; *** = P ≤ 0.001; **** = P ≤ 0.0001. Unpaired t tests were used in panels A-E and the data are shown as mean ± standard deviation. An ordinary one way ANOVA way used for the comparison of m+1 isotopomers (blue) in Panels F-K, while unpaired t tests were used for the comparison of labeling in outer m+1 isotopomers vs inner m+1 istopomers (red). Data are shown as mean ± standard deviation.

A puzzling aspect of in vivo tumor metabolism studies is the presence of large M+1 isotopologue fractions in TCA cycle intermediates after infusion with tracers that produce [1,2-^13^C]acetyl-CoA (e.g. [U-^13^C]glucose). Explanations for these M+1 isotopologues include carboxylation of unlabeled intermediates using ^13^CO_2_ liberated during the oxidation of labeled fuels, and the progressive oxidation of M+2 isotopologues through multiple rounds of the TCA cycle (Duan et al., 2022; Hensley et al., 2016). Simple isotopologue analysis does not discriminate between these possibilities. However, the position of ^13^C within TCA cycle metabolites reflects the mechanism of label delivery to the cycle (Supplementary Fig. 3A,B). Carboxylation reactions label the outer carbons of the product metabolite, and the label is subsequently lost as the product is oxidized. On the other hand, labeling of citrate through condensation of [1,2-^13^C]acetyl-CoA and unlabeled OAA produces M+2 intermediates through the first turn of the cycle, with the ^13^C located at positions 4 and 5 of glutamate and then at either 1 and 2 or 3 and 4 of aspartate. With each subsequent turn, the likelihood of incorporating [1,2-^13^C]acetyl-CoA is proportional to its fractional enrichment, which is low in SK-N-AS tumors (F_C3_=0.3, Fig. 4B). Oxidation of intermediates labeled as M+2 on the first turn results in M+1 labeling, starting in turn 2, but the position of ^13^C is mixed between inner and outer carbons. Specifically, labeling in aspartate under this scenario is predicted to be distributed equally between inner and outer carbons, even with no contribution of ^13^CO_2_ recycling. Examination of M+1 aspartate labeling after natural abundance correction in SK-N-AS xenografts infused with [U-^13^C]glucose revealed nearly equal labeling of outer and inner carbons, with only a non-significant predominance of outer carbon labeling (Fig. 4F). Similar results were observed during infusions with [U-^13^C]acetate, although the overall labeling was lower (Fig. 4G). We also examined aspartate isotopomers in mouse tissues after [U-^13^C]glucose infusion, and again did not detect an excess of outer carbon labeling (Fig. 4H-K). This suggests that TCA cycling rather than carboxylation is the major source of M+1 labeling in aspartate under these conditions.

### Application of LC-MS/MS isotopomer analysis to human cancer

In human cancer studies, we typically use at least 100 mg of tissue to assess ^13^C labeling by NMRS (Courtney et al., 2018; Hensley et al., 2016; Maher et al., 2012; Mashimo et al., 2014). In tumors from the brain and lung, this approach has provided adequate signal to noise ratios (SNR) to observe several isotopomers of glutamate and aspartate. However, using samples this large obscures the regional metabolic heterogeneity characteristic of solid tumors in humans. The problem is compounded in tumors from the kidney, which are both highly heterogeneous and characterized by low labeling of TCA cycle intermediates from [U-^13^C]glucose (Courtney et al., 2018). We compared NMRS and LC-MS/MS analysis of tumors from two kidney cancer patients with hereditary leiomyomatosis and renal cell carcinoma (HLRCC). Both patients were infused with [U-^13^C]glucose during nephrectomy. Kidney tumors in HLRCC lack fumarate hydratase (FH) and are therefore expected to have suppressed TCA cycle function. After nephrectomy, samples were obtained from the nonmalignant kidney and from two sites of the tumor. In one sample from patient A’s tumor, glutamate C4 labeling could not be detected by NMRS (Fig. 5A). In the other sample, weighing 460 mg, NMRS detected ^13^C multiplets in glutamate C3 and C4, but the SNR was low (Fig. 5B). Multiplets at succinate C2/3 were prominent in both samples, perhaps reflecting succinate accumulation resulting from FH loss.

**Figure 5.**
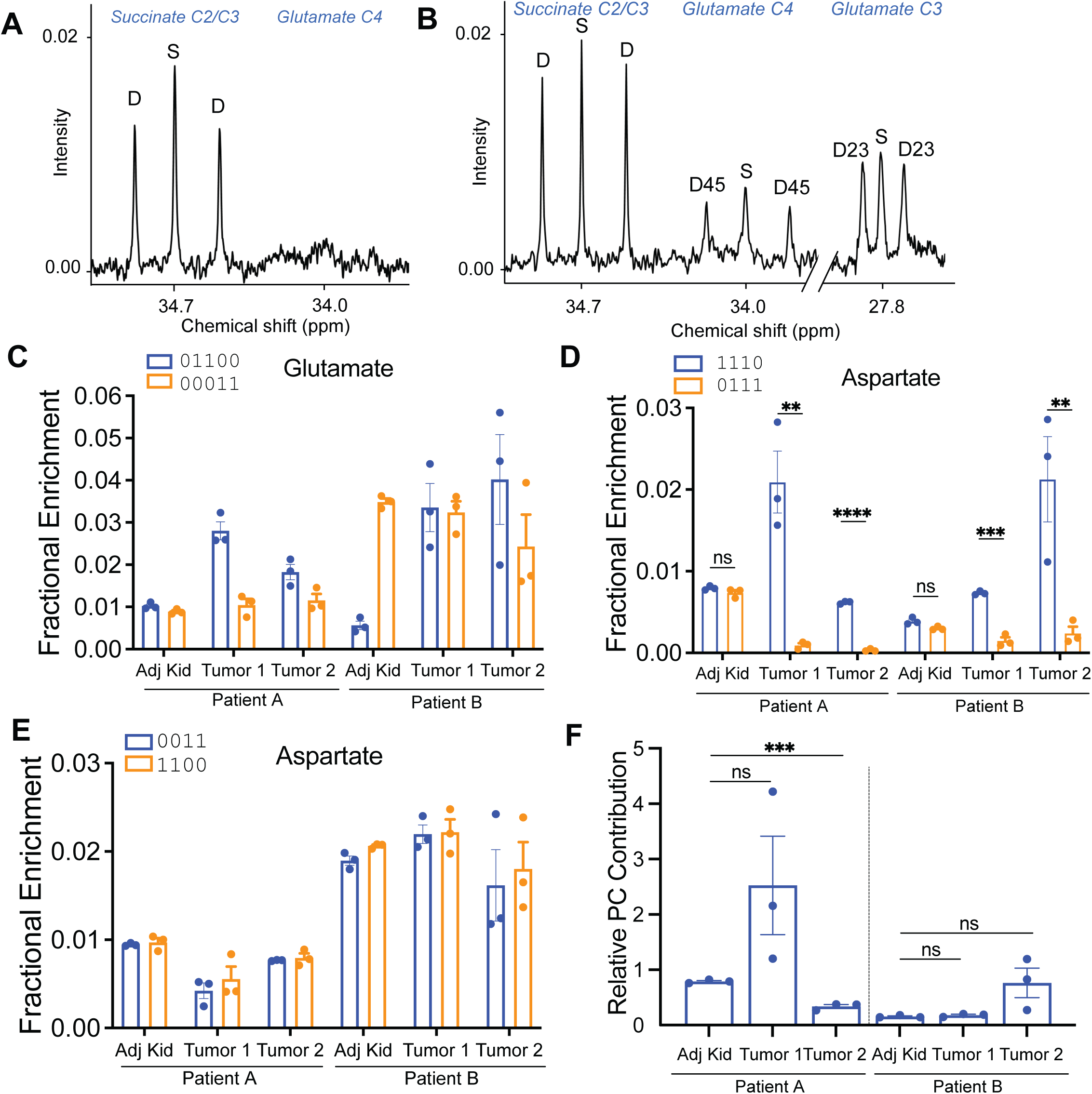
Isotopomer analysis from human kidney tumors. (A-B) Expanded ^13^C NMR spectra of glutamate C4 and C3 from two fragments of a human HLRCC after infusion with [U-^13^C]glucose during nephrectomy. (C) Selected glutamate isotopomers analyzed by LC-MS/MS from fragments of tumor and adjacent, non-malignant kidney (Adj Kid) tissue obtained after infusion with [U-^13^C]glucose during nephrectomy in two patients. (D-E) Relative contributions of selected aspartate M+3 (D) and M+2 (E) isotopomers in these tissue fragments. (F) Relative abundance of aspartate isotopomers derived from PC [(1110+0111)-(1011+1101)], normalized to those derived from PDH (1100+0011). P values: ns = P > 0.05; * =P ≤ 0.05; ** = P ≤ 0.01; *** = P ≤ 0.001; **** = P ≤ 0.0001. Unpaired t tests were used in panel D and F.

LC-MS/MS used three small (10-20 mg) fragments of each sample. Of the observed ion pairs monitored by MRM in these samples, 60-100% had SNRs over 10 (Supplementary Figure 4, Supplementary Data III). Natural abundance corrected glutamate labeling had hallmarks of both PDH and PC activity, with 2-4% of the pool labeled as glutamate 01100 and 1-3% labeled as glutamate 00011 (Fig. 5C). Because analysis of glutamate 01100 is complicated by potential calculation error, we also examined glutamate 111000 proceeding from PC-dependent production of aspartate 0111. However, the abundance of this isotopomer was negligible. Interestingly, aspartate 1110 exceeds aspartate 0111 in both fragments of the tumor (Fig. 5D). These patterns imply activity of PC to label OAA/aspartate, but poor equilibration with the fumarate pool, as predicted by loss of FH activity within these tumors (Supplementary Figure 5A,B). In the nonmalignant kidney fragments from both patients, where FH activity persists, aspartate 1110 and 0111 are equivalent (Fig. 5D), indicating that the lack of symmetrization in the tumors was not an artifact of tissue handling during the surgery. It is also interesting that aspartate isotopomers arising from PDH (1100 and 0011) were equivalent to each other in both the tumors and the kidney samples (Fig. 5E), indicating that the expected symmetry is achieved in isotopomers arising from progression of the cycle from citrate to OAA/aspartate. It is unclear whether the equivalent levels of aspartate 1100 and 0011 in the tumors reflects a small amount of residual FH activity in tumor cells, or whether part of the aspartate pool in these samples was taken up from other tissues with functional FH.

Finally, to estimate the relative contributions of PC and PDH activity to TCA cycle labeling, we determined the relative abundance of aspartate isotopomers derived from PC (specifically, the [(1110+0111)-(1011+1101)] difference discussed above) and derived from PDH (1100+0011). This ratio was below 1.0 in the non-malignant kidneys and highly variable among the tumor samples, with ratios in some fragments exceeding 2.0 (Fig. 5F). This analysis reveals the ability of the LC-MS/MS method to detect regional metabolic heterogeneity from small samples of primary human tumors and to provide information about relative routes of carbon entry into and processing by the TCA cycle.

## DISCUSSION

Although ^13^C tracers are excellent tools to assess metabolic activity, simple isotopologue analysis (by far the most common mass spectrometry technique currently used in such experiments) reports only a fraction of the information encoded by ^13^C labeling (Buescher et al., 2015). We describe a method that combines the high sensitivity of mass spectrometry with the positional information usually determined by NMRS. A primary objective was to be able to report the isotopomers of glutamate required in calculations of metabolic parameters relevant to the TCA cycle, including anaplerosis and enrichment of the acetyl-CoA pool that supplies citrate synthesis. In addition to this subset of glutamate isotopomers, the method provides direct or indirect information about the complete set of isotopomers from both glutamate and aspartate. The complementary information provided by all these labeled species provides a highly detailed analysis of the TCA cycle, while requiring far less material than what is needed for NMRS (only 1% in experiments using cultured cells).

The method also has a few limitations worth considering. First, although the information yield is potentially even greater than NMRS, the technique is analytically demanding compared to NMRS. For straightforward applications (e.g. calculation of F_C3_) when the SNR is high and tissue is abundant, NMRS may still be preferable. Second, due to the chemical nature of glutamate and aspartate, not every desired fragment can be generated. As a result, the current method still requires some simulation and results in nontrivial errors (up to 3%) for some isotopomers. This limitation can be mitigated somewhat by choosing labeling strategies that minimize the contributions of isotopomers with relatively high levels of uncertainty, and by capitalizing on complementary pieces of information (e.g. aspartate isotopomers and F_C3_) that reduce the dependence on uncertain glutamate isotopomers.

We believe this approach will have particular value in human and other in vivo studies when performing multiple tracer experiments is impractical or impossible. For example, specialized tracers can be used to improve certainty about specific reactions in the metabolic network. The classical use of [3,4-^13^C]glucose to probe PC activity is one example. However the high cost and otherwise low information yield of this tracer makes it less appealing than [U-^13^C]glucose in human studies. The isotopomer method’s increased clarity about PC activity downstream of [U-^13^C]glucose is therefore a considerable advantage. The high sensitivity of the method is also an obvious advantage for human cancer studies, where metabolite labeling tends to be low owing to the sub-maximal enrichment of the precursor pool in the circulation. Although it is possible to obtain informative 1D ^13^C NMR spectra from tumors, the large samples required make it difficult to study regional heterogeneity of ^13^C signatures, which can be substantial in solid tumors (Hensley et al., 2016). The high sensitivity of the method described here should make it possible to perform isotopomer analysis in very small tumor samples or perhaps even populations of distinct cell types from the tumor microenvironment.

Finally, a challenge of in vivo isotope tracing studies in cancer is the appearance of unexpected isotopologue distributions. Verifying the origin of such labeling patterns is straightforward in cultured cells, where silencing enzymes of interest makes it possible to identify the responsible pathway. This is more difficult in mice and impossible in patients. The prominent M+1 labeling of TCA cycle intermediates after infusions with uniformly ^13^C-labeled nutrients in vivo is an example of this challenge. While M+1 fractions tend to be small in cell culture, this fraction appears quickly in vivo and matches or exceeds the more familiar M+2 fraction. Positional specificity is informative about the origins of the label. If ^13^CO_2_ liberated from labeled substrates is used as substrate in carboxylation reactions, it is predicted to reside on the outer positions (i.e. carbons 1 and 4) of 4-carbon intermediates like aspartate. These labels are largely lost as ^13^CO_2_ during subsequent decarboxylation reactions rather than transferred to internal carbons. On the other hand, PDH followed by successive turns of the cycle produces labels distributed evenly between internal and external carbons. Simulations of TCA cycle activity indicate that when F_C3_ is low, these successive rounds of the cycle can produce high levels of M+1 isotopologues, sometimes exceeding the M+2 fraction, even in the absence of ^13^CO_2_ fixation (Hensley et al., 2016). We detect nearly identical labeling of internal and external aspartate carbons after infusion with [U-^13^C]glucose or [U-^13^C]acetate. This suggests that most of the M+1 labeling in TCA cycle intermediates arises downstream of PDH rather than carboxylation and ^13^CO_2_ recycling. We emphasize that the data do not rule out some contribution of ^13^CO_2_ recycling, but the simplest interpretation is that this is a minor component of the M+1 labeling observed in vivo.

In summary, we established an LC-MS/MS method to identify complete isotopomer fractions of glutamate or aspartate. The isotopomer information acquired using this method can detect PDH activity, PC activity, anaplerosis and somatic loss of FH in tumors. These examples demonstrate the utility of this approach in tumor metabolism and other biological systems.

## Supporting information

Supplemental Data III

## ACKNOWLEDGEMENTS

This article is subject to HHMI’s Open Access to Publications policy. HHMI lab heads have previously granted a nonexclusive CC BY 4.0 license to the public and a sublicensable license to HHMI in their research articles. Pursuant to those licenses, the author-accepted manuscript of this article can be made freely available under a CC BY4.0 license immediately upon publication. This research was supported by the Howard Hughes Medical Institute Investigator’s Program (R.J.D) and grants from the N.I.H. (R35CA220449, P50CA196516 and 2P50CA070907 to R.J.D.; P41-122698, 5U2CDK119889, and NIH R01-132254 to M.E.M), the National Science Foundation (NSF, DMR-1644779 to M.E.M), and the Cancer Prevention and Research Institute of Texas (CPRIT, RP180778 to R.J.D). D.B was supported by NCI F31CA239330. A portion of this work was performed at the National High Magnetic Field Laboratory, which is supported by National Science Foundation Cooperative Agreement number DMR-1644779, & the State of Florida. The content of this manuscript is solely the responsibility of the authors and does not necessarily represent the official views of the NIH.

## AUTHOR CONTRIBUTIONS

Conceptualization: F.C., R.J.D.; Investigation: F.C., D.B., L.C., R. M., Z. W., P.P., C. Y., S. K., W. G., H.S.V., T.M., L.Z., M.M.S., D. D., K. C.O., E.J., V.M., C.R.M, and M.E.M.; Writing – Original Draft: F.C., D.B., R.J.D.; Writing – Reviewing and Editing: F.C., D.B., L.C., R. M., Z. W., P.P., C. Y., S. K., W. G., H.S.V., T.M., L.Z., M.M.S., D. D., K. C.O., E.J., V.M., C.R.M, M.E.M, R.J.D.; Visualization: F.C., D.B., R.J.D.; Funding Acquisition: R.J.D.; Resources: V.M., C.R.M., M.E.M., R.J.D.; Supervision: R.J.D.

## METHODS

### Lead Contact

Further information and requests for resources and reagents should be directed to the lead contact, Ralph DeBerardinis, MD, PhD.

### Materials Availability

This study did not generate new reagents.

### Data and Code Availability

The script for natural isotope abundance correction matrix, non-negative least square regression, and error estimation have been deposited at the Github repository (https://github.com/RJDlab/Glu_Asp_Isotopomers).

### Experimental Model and Human Subject Details

Human subjects research was approved by the Institutional Review Board of UT Southwestern Medical Center (“An Investigation of Tumor Metabolism in Patients Undergoing Surgical Resection” (STU062010-157)). Patients were selected for inclusion based on imaging and clinical features consistent with renal cell carcinoma.

### Chemicals

All chemicals and reagents were LC-MS grade or higher. Sterile, pyrogen free [U-^13^C]glucose was purchased from Cambridge Isotope Laboratories for human infusions (CLM-1396-MPT-PK). For mouse infusions, [U-^13^C]acetate was purchased from Sigma-Aldrich (663859) and [U-^13^C]glucose was purchased from Cambridge Isotope Laboratories (CLM-1396).

### Human Infusions

Sterile, pyrogen free [U-^13^C]glucose was infused at the time of nephrectomy. A peripheral intravenous line was placed on the morning of surgery and labeled glucose was infused as a bolus of 8g over 10 minutes followed by 8g per hour as a continuous infusion. Standard procedures were followed for tumor resection and tissue fragments were flash frozen in liquid nitrogen. All diagnoses were made by the attending clinical pathologist. These infusion parameters are consistent with previously published work (Faubert et al 2021, Courtney et al 2018, Faubert et al., 2017; Hensley et al., 2016; Maher et al., 2012; Mashimo et al., 2014).

### Mouse Infusions

All mouse infusions were performed in compliance with protocols approved by the Institutional Animal Care and Use Committee at the University of Texas Southwestern Medical Center (Protocol 2016-101694). For tumor studies, subcutaneous injections were done in the right flank of NOD.CB17-Prkdc*^scid^*Il2rg^tm1Wjl^/SzJ (NSG) male and female mice aged 4-8 weeks old. To establish xenografts from cancer cell lines, suspensions of neuroblastoma cells were prepared for injection in RPMI 1640 media with Matrigel (CB-40234; Fisher Scientific). 50μL of the cell suspension was combined with 50μL Matrigel for a total volume of 100μL per mouse. For studies that involved treatment with the complex I inhibitor (IACS-010759, ChemieTek), the mice were administered IACS-010759 by oral gavage every day for 5 days (10 mg/kg body mass in 100 μL of 0.5% methylcellulose and 4% DMSO) once subcutaneous tumors reached 200mm^3^. On the 5^th^ day, mice were infused with [U-^13^C]glucose or [U-^13^C]acetate and tumors were harvested 3-5 hours following the last treatment dose, depending on the infusion. Mice were weighed and subcutaneous tumors measured at the beginning and end of the 5-day treatment period. Mice were anesthetized and then a 27-gauge catheter was placed in the lateral tail vein under anesthesia. [U-^13^C]acetate (Sigma-Aldrich, 663859) was delivered as a bolus of 0.3 mg/g body mass over 1 min in 150 μL of saline, followed by continuous infusion of 0.0069 mg/g body mass/min for 3 hours in a volume of 150 μL/hour. [U-^13^C]glucose (Cambridge Isotope Laboratories, CLM-1396) was intravenously infused as a bolus of 0.4125 mg/g body mass over 1 min in 125 μL of saline, followed by continuous infusion of 0.008 mg/g body mass/min for 3 hours in a volume of 150 μL/hour. At the end of the infusion, mice were euthanized and tumors and/or organs were harvested and immediately frozen in liquid nitrogen. To assess the fractional enrichments in plasma, 20 μL of blood was obtained after 30, 60, 120, and 180 minutes of infusion.

### Cell lines and culture conditions

H460 cells were obtained from the Hamon Cancer Center Collection (University of Texas Southwestern Medical Center) and maintained in RPMI-1640 supplemented with penicillin–streptomycin and 5% fetal bovine serum (FBS) at 37°C in a humidified atmosphere containing 5% CO_2_. SF188-derived glioblastoma cells overexpressing human Bcl-xL (SFxL) were previously reported (Wise et al., 2008) and Huh-7 hepatocellular carcinoma cells were a gift from Professors Michael S. Brown and Joseph L. Goldstein. SFxL and Huh-7 were maintained in Dulbecco’s modified Eagle medium (DMEM) supplemented with penicillin–streptomycin and 10% FBS at 37°C in a humidified atmosphere containing 5% CO_2_. All cell lines were validated by DNA fingerprinting using the PowerPlex 1.2 kit (Promega) and were confirmed to be mycoplasma-free using the e-Myco kit (Boca Scientific).

### PC depletion in Huh7 cells

lentiCRISPR v2 was a gift from Feng Zhang (Addgene plasmid # 52961; http://n2t.net/addgene:52961; RRID:Addgene_52961). Guide RNA oligos were ordered from IDT Company. Guide RNAs (sgScr: 5’ to 3’ TTCTTAGAAGTTGCTCCACG, sgPC#1: 5’ to 3’ CAGGCCGCGGCCGATGAGAT, sgPC#2: 5’ to 3’ ACAGGTGTTCCCGTTGTCCC) were cloned into LentiCRISPRv2 (Sanjana et al., 2014), then transfected into 293T cells using Lipofectamine 3000 (Thermo Fisher Science, L3000015) with a 2:1 ratio of psPAX2: pMD2G. Medium containing viral particles was collected and filtered using 0.45µM membrane filters 48 hours after the transfection. Huh7 cells were cultured in media containing lentivirus and 4µg/mL polybrene (Sigma, TR-1003-G) for 24 hours followed by selection in 5µg/mL puromycin until the uninfected control cells died.

### Western Blots

Cells were lysed in RIPA buffer (Boston BioProducts, BP-115) containing protease and phosphatase inhibitors (Thermo Fisher Sciences, 78444), then centrifuged at 4°C for 10 minutes at ∼20,160 g. Supernatants were transferred to new pre-chilled 1.5 mL tubes and protein concentrations were quantified using the Thermo Fisher Pierce BCA Assay Kit (Thermo Fisher, 23225). Protein lysates were resolved via SDS-PAGE and transferred to PVDF membranes. Membranes was blocked in 5% bovine serum albumin (BSA) in Tris Buffered Saline with Tween-20 (TBST (20 mM Tris pH 7.5, 150 mM NaCl, 0.1% Tween-20)) and then incubated with primary antibodies (PC: Proteintech, 16588-1-AP, 1:1000 dilution, or β-actin: Cell Signaling, 8457S, 1:1000 dilution) in TBST supplemented with 5% BSA at 4°C overnight. Primary antibodies were detected with a horseradish peroxidase-conjugated secondary antibody (Cell Signaling Technology, 7074S, 1:2000 dilution) for 1 hour followed by exposure to ECL reagents (Fisher Scientific, PI32106).

### Sample preparation for LC-MS/MS

Tissue samples (2-5mg) were cut on a surface cooled with dry ice, transferred into a 1 mL microcentrifuge tube on dry ice, and ground with a pellet pestle. Ice cold 80% (vol/vol) acetonitrile in water (200 µL) was added to the tube, and the tissue was homogenized until no visible chunks remained.

Cells were plated at 1 × 10^5^ cells per 6-cm plate 16 h before labelling. The next day, cells were incubated in glucose/glutamine-free DMEM 20 mM [U-^13^C]glucose or [3,4-^13^C]glucose. At the desired time, the medium was aspirated and cells were rinsed with ice cold saline. The saline was aspirated, 80% (vol/vol) acetonitrile in water was added and the cells were collected with a cell scraper. The resulting mixture was transferred into a microcentrifuge tube and subjected to three freeze-thaw cycles between liquid nitrogen and a 37 °C water bath. The samples were vortexed for 1 min before centrifugation at ∼20,160 g for 15 min at 4°C. The metabolite-containing supernatant (195 µL) was transferred into pre-cut Bond® Elut PH (100 mg, 1 ml) SPE cartridges (Agilent, 12102005) placed in 1.7 mL microcentrifuge tubes and then centrifuged at ∼3000 g for 3 min. Alternatively, the supernatants were passed through Oasis® HLB LP 96-well plate 60µm (60mg) SPE cartridges (Waters, 186000679). The filtrate was then transferred to LC/MS vials for analysis.

### LC-MS/MS analysis

Samples were analyzed on an AB Sciex 6500 QTRAP liquid chromatography/mass spectrometer (Applied Biosystems SCIEX) equipped with a vacuum degasser, quaternary pump, autosampler, thermostatted column compartment and triple quadrupole/ion trap mass spectrometer with electrospray ionization interface, and controlled by AB Sciex Analyst 1.6.1 Software. SeQuant® ZIC®-pHILIC 5µm polymer (150mm×2.1mm) columns were used for separation. Solvents for the mobile phase were 10 mM ammonium acetate aqueous (pH 9.8 adjusted with NH_3_.H_2_O (A) and pure acetonitrile (B). The gradient elution was: 0–20 min, linear gradient 90-65% B, 20–23 min, linear gradient 65-30% B, 23-28 min, 30% B, and 28–30 min, linear gradient 30-90% B then reconditioning the column with 90% B for 5 min. The flow-rate was 0.2 ml/min and the column was operated at 40°C.

### 13C Nuclear Magnetic Resonance Spectroscopy

For the cell culture, experiments, H460 cells were plated at 5 × 10^6^ cells per 15 cm plate in three plates and allowed to adhere for 16 h before labelling. The next day, cells were incubated in glucose/glutamine-free DMEM media containing 20 mM [U-^13^C]glucose for 24h before collection. The medium was aspirated and cells were rinsed with ice cold saline. The saline was aspirated and 80% (vol/vol) acetonitrile in water (5mL) was added to the cells. The cells were scraped with a cell scraper. The mixtures were transferred into microcentrifuge tubes (0.5ml each) and subjected to three freeze-thaw cycles between liquid nitrogen and a 37°C water bath. The samples were vortexed for 1 min before centrifugation at ∼20,160 g for 15 min at 4°C and the supernatants were centrifuged again at ∼20,160 g for 15 min at 4°C before being dried at room temperature in a Speedvac system (Thermo Scientific, Waltham, MA).

The sample for ^13^C NMR analysis was prepared by dissolving the dried residue in 54μl of 50mM sodium phosphate buffer in D_2_O containing 2mM ethylenediaminetetraacetic acid (EDTA). A 6 μl solution of 0.5 mM deuterated sodium 3– trimethylsilyl–1–propane sulphonate (d_6_-DSS) (internal standard) and 0.2% sodium azide (NaN_3_) in D_2_O was added to the sample. The solution was vortexed thoroughly, centrifuged at 10,000 g for 15 minutes and supernatant was loaded into a 1.5 mm NMR tube. ^13^C NMR data were acquired using a 14.1 T NMR magnet equipped with a ^13^C-optimized home-built high temperature superconducting (HTS) probe (Ramaswamy et al., 2013) and VNMRJ Version-4.0 software. The following parameters were used to acquire ^13^C NMR data: number of scans = 30412, acquisition time (AQ) = 1.5s, relaxation delay (d1) = 1.5s, flip angle = 45°, acquired size = 54k. The ^13^C NMR spectrum was Fourier Transformed with an exponential line-broadening factor of 0.5 Hz, zero filling to 128k data points, and applying Whittaker smother baseline correction in MestReNova v14.0.1-23284 (Mestrelab Research S.L.). Mixed Gaussian/Lorentzian shape was used in the line fitting tool to fit the peak area of the multiplets for each of the ^13^C NMR peaks of glutamate and aspartate.

For NMR of samples from HLRCC patients, frozen tissue samples (0.45 - 0.48 g) were powdered using a mortar and pestle chilled with liquid nitrogen. Ground tissue was transferred into a 15-mL conical tube containing perchloric acid solution (10%; 3 mL) on ice, vortexed for 1 min, and centrifuged at 25,000 g and 4°C for 10 min. The supernatant was transferred to a new 15-mL conical tube, and the extraction was repeated by adding perchloric acid solution to the pellet. The combined supernatant was neutralized with KOH, centrifuged, and the supernatant was transferred to a 20-mL glass vial. After drying under vacuum, the extracts were dissolved in heavy water (^2^H_2_O, 200 μL) containing 4,4-dimethyl-4-silapentane-1-sulfonic acid (5 mM), centrifuged at 20,000 g for 5 min and the supernatant was transferred to a 3-mm NMR tube.

NMR spectra were collected using a 14.1T spectrometer (Varian INOVA, Agilent, Santa Clara, CA) equipped with a 3-mm broadband probe with the observe coil tuned to ^13^C (150 MHz). NMR spectra were acquired using a 45° pulse, a 36,765-Hz sweep width, 55,147 data points, and a 1.5-sec acquisition time with 1.5-s interpulse delay at 25°C. Proton decoupling was performed using a standard WALTZ-16 pulse sequence. Spectra were averaged with 25,000 scans and a line broadening of 0.5 Hz was applied prior to Fourier transformation. Spectra were analyzed using ACD/Labs NMR spectral analysis program (Advanced Chemistry Development, Inc., Toronto, Canada).

### Nonnegative least square regression

Nonnegative least square regression was implemented using the glmnet R package (Friedman et al., 2010), with the following parameters: lambda = 0, lower.limits = 0, intercept = FALSE and thresh = 1e-30, such that the regression is not regularized, the coefficients are bounded between 0 and 1, and the convergence threshold is small but can generally be reached within the default number of iterations. We compared this method to other R packages with Lawson-Hanson’s algorithm (Lawson et al., 1995) implementation such as in the NNLS (Katharine M. Mullen, 2012) or pracma (Borchers, 2021) R packages and found the glmnet-based method to provide the lowest error from our simulation error estimation (Supplementary materials Fig S-I 1 and 2).

### Natural Abundance Correction

The following isotopes were considered during natural abundance correction: ^2^H (0.0156%), ^13^C (1.082%), ^15^N (0.366%), ^17^O (0.038%), and ^18^O (0.204%). All possible ^2^H, ^13^C, ^15^N, ^17^O, and ^18^O isotopomers of the precursor and product ions are listed and their probabilities are calculated and assigned to each MRM transition. The calculation matrices of glutamate and aspartate with or without natural isotope abundance correction are available in Supplementary Data II. The R script used for calculating the correction matrix can be found at the GitHub repository (https://github.com/RJDlab/Glu_Asp_Isotopomers).

## STATISTICAL ANALYSIS

Samples were analyzed as described in the figure legends. Data were considered significant if p<0.05. Statistics were calculated using PRISM software, and statistical details can be found in the figure legends for each figure.

---

**Figure S1.**
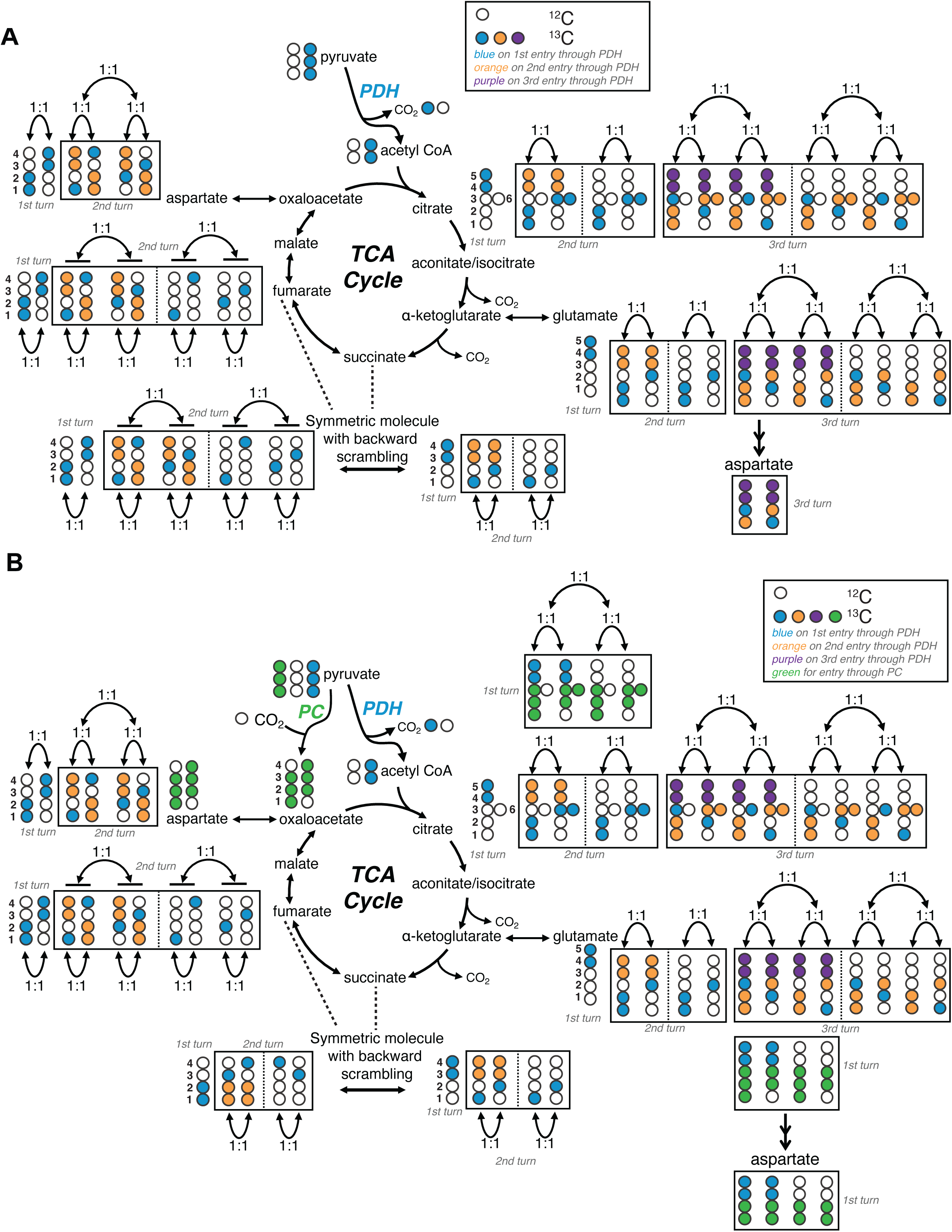
Schematic of isotopomer labeling from [U-^13^C]pyruvate. **A).** Isotopomer schematic where PDH is active and PC is inactive. White circles indicate ^12^C, while colored circles indicate ^13^C. Succinate and fumarate generate mirrored isotopomers due to the symmetric nature of these metabolites. These mirrored isotopomers give rise to corresponding isotopomers in subsequent steps of the cycle, ultimately producing symmetric isotopomer pairs in aspartate with equal distribution in 1000/0001, 0100/0010, 1100/0011, and 1110/0111/1011/1101. Aspartate-1111 can arise from glutamate 11111 or 01111. The distribution of aspartate m+1 is discussed in Supplemental Figure 3A. **B)** Isotopomer schematic where both PDH and PC are active. PC produces a new isotopomer of oxaloacetate (1110, green), which can then be converted to 0111 after equilibration with symmetric TCA cycle intermediates and eventually producing malate, oxaloacetate and aspartate 1110/0111. The large impact of PC on 1110 and 0111 means that these isotopomers can exceed 1011 and 11011; note that in Panel A, where PC is inactive, 1110, 0111, 1011 and 1101 are all present at equivalent abundances. Therefore the excess of 1110 and 0111 indicates the presence of PC activity.

**Figure S2.**
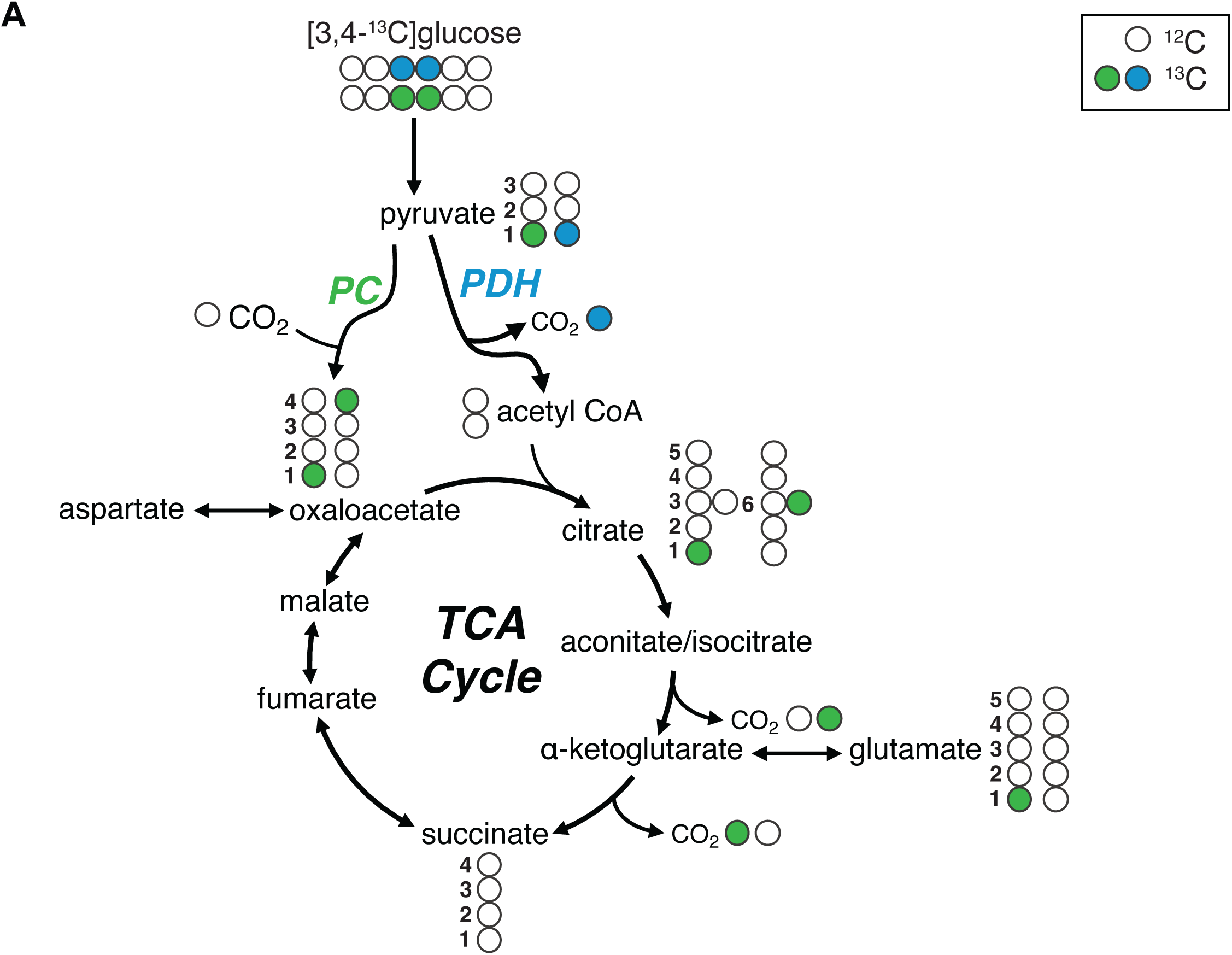
Schematic of isotopomer labeling from [3,4-^13^C]glucose. [3,4-^13^C]glucose is converted to [1-^13^C]pyruvate through glycolysis and enters into the TCA cycle through PDH or PC. If [1-^13^C]pyruvate enters the TCA cycle through PDH, the ^13^C at the C1 position will become ^13^CO_2_, resulting in unlabeled acetyl-CoA. However, if [1-^13^C]pyruvate enters the TCA cycle through PC, then the labeled carbon at C1 is retained as [1-^13^C]oxaloacetate and later [4-^13^C]oxaloacetate through symmetrization. [1-^13^C] and [4-^13^C]oxaloacetate become [1-^13^C] and [6-^13^C]citrate, and eventually [1-^13^C]α-ketoglutarate and [1-^13^C]glutamate. When [1-^13^C]α-ketoglutarate is converted to succinate via succinyl-CoA, the remaining ^13^C is released as ^13^CO_2_ and subsequently produces unlabeled four carbon intermediates.

**Figure S3.**
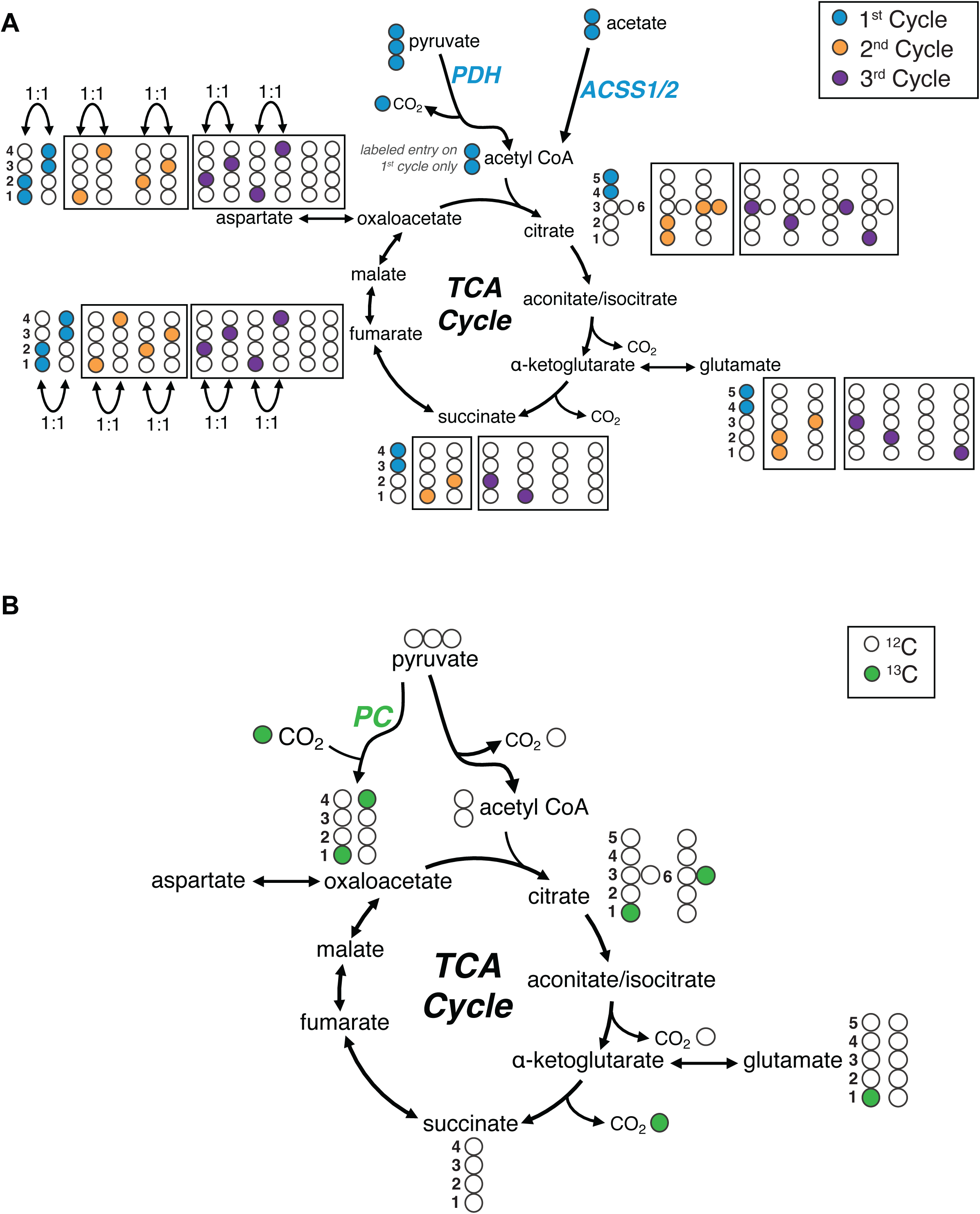
Schematic of M+1 isotopomer labeling. **A)** Isotopomer schematic where PDH is active and PC is inactive. Equal distribution of both interior isotopomers (0100/0010) and exterior isotopomers (1000/0001) are generated in aspartate and other four-carbon intermediates due to backward scrambling in fumarate. [1,2-^13^C]acetyl-CoA is shown entering only on the first cycle for simplicity, and can arise from [U-^13^C]pyruvate or [U-^13^C]acetate. **B)** Isotopomer schematic where both PDH and PC are active with an unlabeled source of pyruvate. ^13^CO_2_ enters the TCA cycle through PC and it is converted to [4-^13^C]oxaloacetate and [1-^13^C]oxaloacetate, the latter arising through fumarate/succinate scrambling. [1-^13^C] and [4-^13^C]oxaloacetate will become [1-^13^C] and [6-^13^C]citrate, and eventually [1-^13^C]α-ketoglutarate and [1-^13^C]glutamate. When [1-^13^C]α-ketoglutarate is converted to succinate via succinyl-CoA, the remaining ^13^C is released as ^13^CO_2_ and subsequently produces unlabeled four-carbon intermediates. Therefore, only [1-^13^C] and [4-^13^C]oxaloacetate (and [1-^13^C] and [4-^13^C]aspartate) are generated when the labeled carbon source is ^13^CO_2_ incorporation from PC. Note that other carboxylase reactions giving rise to TCA cycle intermediates also result in labeling of external rather than internal carbons.

**Figure S4.**
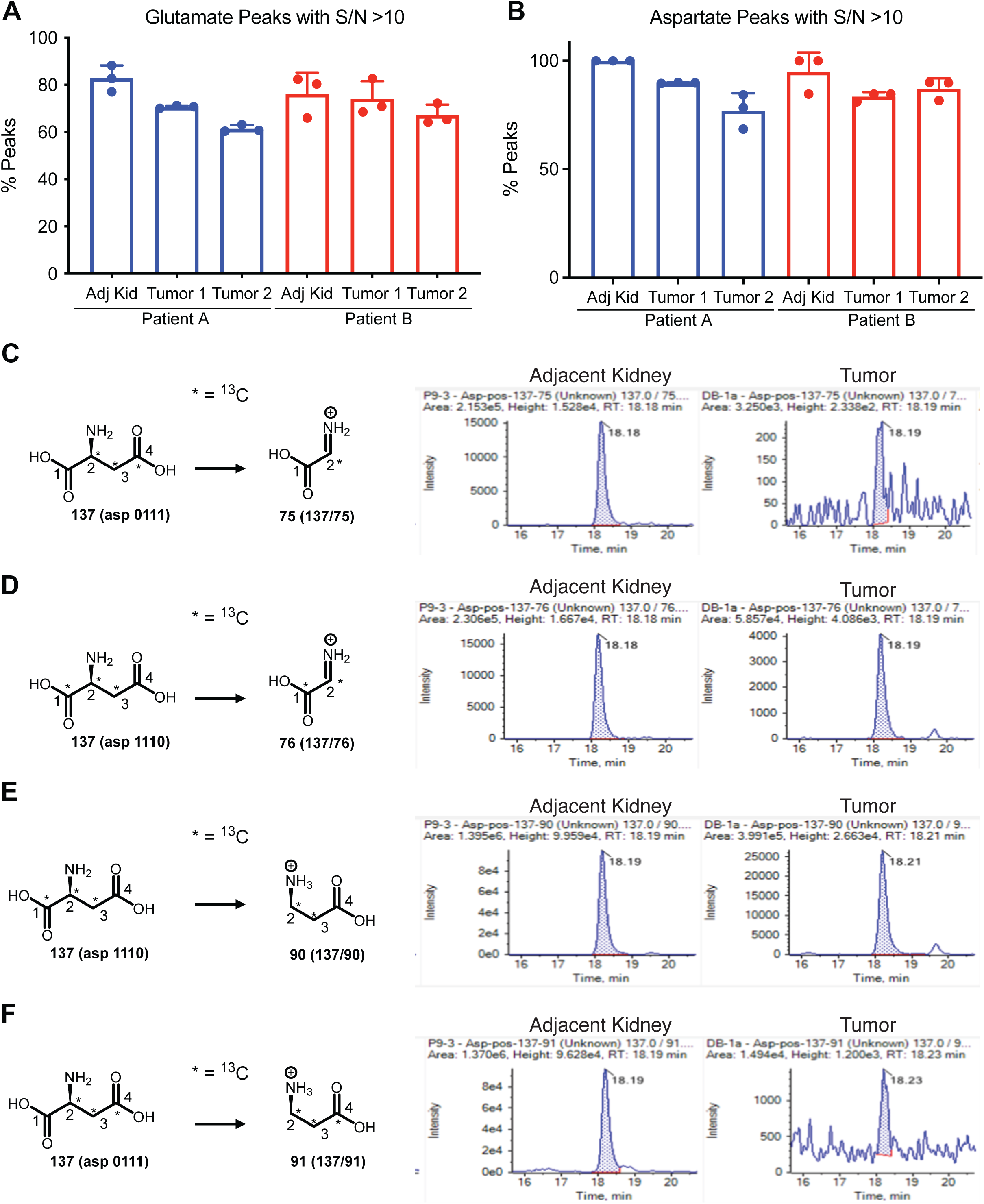
Sensitivity of LC-MS/MS. A-B) Summary of signal to noise ratios for HLRCC patient samples in glutamate and aspartate, respectively. C-F) Representative chromatograms of peaks related to aspartate 1110 in adjacent kidney and tumor samples. Panels C and D reflect ion pairs that report labeling at C1 and C2 (137/75 and 137/76), which provide information about aspartate 0111 and 1110, respectively. Panels E and F reflect ion pairs that report labeling at C2, C3, and C4 (137/90 and 137/91), which provide information about aspartate 1110 and 0111, respectively.

**Figure S5.**
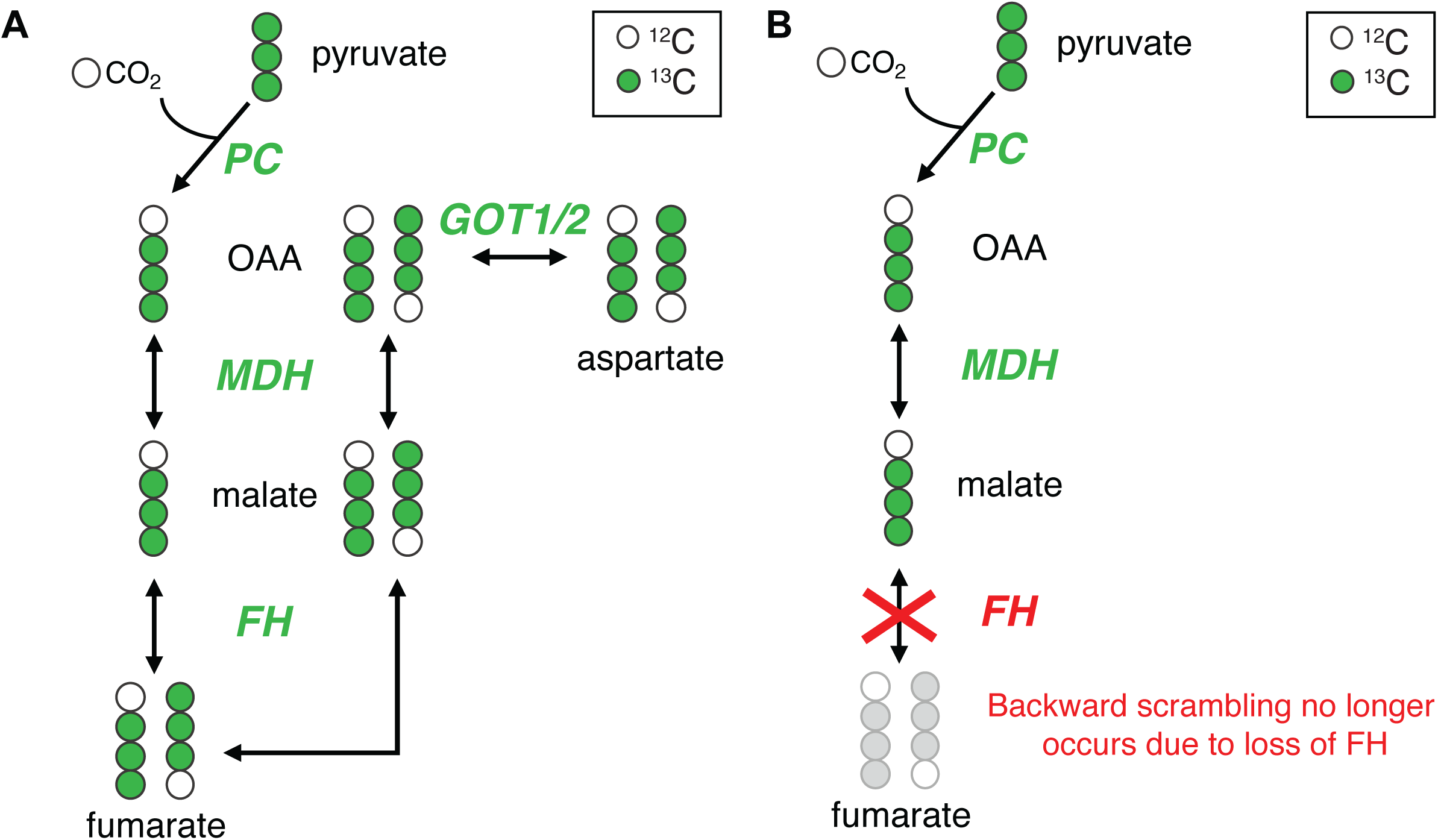
Backward scrambling of succinate and fumarate. A) Schematic of symmetric isotopomers of malate, fumarate, and aspartate arising from equilibration of [1,2,3-^13^C]oxaloacetate with malate and fumarate via malate dehydrogenase (MDH) and fumarate hydratase (FH). The activity of glutamic-oxaloacetatic transaminases (GOT1/2) ultimately produces equivalent abundances of [1,2,3-^13^C]aspartate and [2,3,4-^13^C]aspartate. **B)** When FH is defective, equilibration of symmetric isotopomers does not occur, resulting in an excess of [1,2,3-^13^C]aspartate relative to [2,3,4-^13^C]aspartate (i.e. more aspartate 1110 than 0111).

