## Supplemental Data I for "Comprehensive isotopomer analysis of glutamate and aspartate in small tissue samples"

### MRM methods for glutamate and aspartate isotopomer analysis

The MRM method for glutamate and aspartate isotopomer analysis is described herein.

#### Selected precursor/product ion pairs with mixed fragments

The schemes of glutamate and aspartate fragmentation are shown in Figure 1A and 1B. The fragments of 146/102, 148/56, 146/41 of glutamate and the fragment of 132/88 of aspartate are composed of both a major fragment and a minor fragment. Two additional fragments for glutamate are shown below.

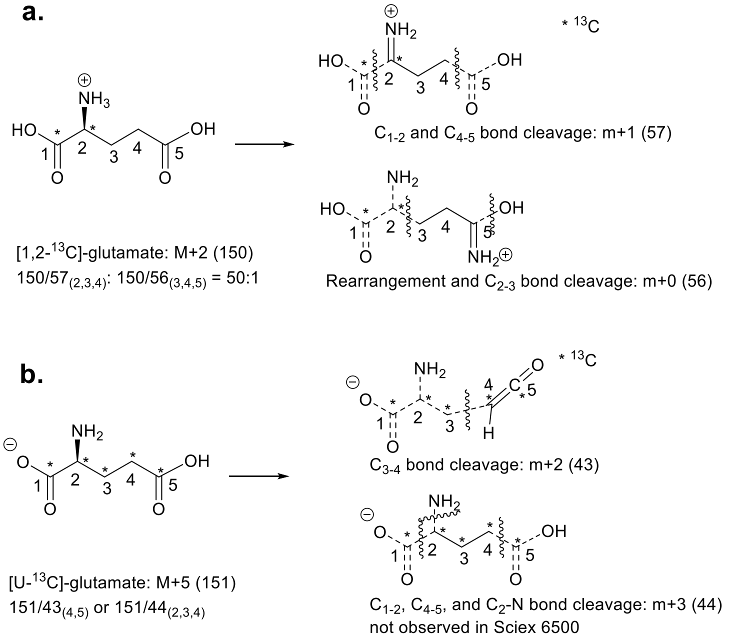


**Scheme 1 Mixed Fragments of Glutamate**. a. Mixed fragments both generating 148/56. b. Mixed fragments both generating 146/41.

The ion pair information for glutamate is listed in Table S1 and the ion pair information for aspartate is listed in Table S2 in the Supplementary Data II file. The actual collision energy (CE) used is modified according to the dynamic range of the mass detector and the optical CE is also provided for reference.

#### Normalized ion pair fractions

For any ion pair whose signal is not detected, a pseudo value of 100 is assigned to eliminate zero values for normalization. The raw data are ion abundance (integration of the area under the curve) of each ion pair, and they are normalized to the total ion abundance.

Step 1: Ion abundance:

| | |
|---|---|
| glutamate_pos-148.1-56.1_2-4_0/0 | 119333971.2 |
| glutamate_pos-149.1-56.1_2-4_1/0 | 4472925.743 |
| glutamate_pos-149.1-57.1_2-4_1/1 | 10326573.24 |
| glutamate_pos-150.1-56.1_2-4_2/0 | 1227541.389 |
| glutamate_pos-150.1-57.1_2-4_2/1 | 83830870.33 |
| glutamate_pos-150.1-58.1_2-4_2/2 | 6276976.271 |
| glutamate_pos-151.1-56.1_2-4_3/0 | 51724.61316 |
| glutamate_pos-151.1-57.1_2-4_3/1 | 3160726.511 |
| glutamate_pos-151.1-58.1_2-4_3/2 | 31346120.89 |
| glutamate_pos-151.1-59.1_2-4_3/3 | 1466476.896 |
| glutamate_pos-152.1-56.1_2-4_4/0 | 10106.61115 |
| glutamate_pos-152.1-57.1_2-4_4/1 | 627915.9406 |
| glutamate_pos-152.1-58.1_2-4_4/2 | 35284349.78 |
| glutamate_pos-152.1-59.1_2-4_4/3 | 16721837.02 |
| glutamate_pos-153.1-56.1_2-4_5/0 | 18819.28065 |
| glutamate_pos-153.1-57.1_2-4_5/1 | 33035.71976 |
| glutamate_pos-153.1-58.1_2-4_5/2 | 311855.5924 |
| glutamate_pos-153.1-59.1_2-4_5/3 | 32220066.8 |

Step 2: Total ion abundance for ion pairs associated with 148/56:

| | |
|---|---|
| Total ion abundance of ion pairs associated with 148/56 | 346721893.8 |

Step 3: Normalization: Each ion abundance is divided by the total ion abundance.

| | |
|---|---|
| glutamate_pos-148.1-56.1_2-4_0/0_normalized fraction | 0.344177779 |
| glutamate_pos-149.1-56.1_2-4_1/0_normalized fraction | 0.012900615 |
| glutamate_pos-149.1-57.1_2-4_1/1_normalized fraction | 0.029783447 |
| glutamate_pos-150.1-56.1_2-4_2/0_normalized fraction | 0.003540421 |

| | |
|---|---|
| glutamate_pos-150.1-57.1_2-4_2/1_normalized fraction | 0.2417813 |
| glutamate_pos-150.1-58.1_2-4_2/2_normalized fraction | 0.018103778 |
| glutamate_pos-151.1-56.1_2-4_3/0_normalized fraction | 0.000149182 |
| glutamate_pos-151.1-57.1_2-4_3/1_normalized fraction | 0.009116028 |
| glutamate_pos-151.1-58.1_2-4_3/2_normalized fraction | 0.0904071 |
| glutamate_pos-151.1-59.1_2-4_3/3_normalized fraction | 0.004229548 |
| glutamate_pos-152.1-56.1_2-4_4/0_normalized fraction | 2.9149E-05 |
| glutamate_pos-152.1-57.1_2-4_4/1_normalized fraction | 0.001811007 |
| glutamate_pos-152.1-58.1_2-4_4/2_normalized fraction | 0.101765566 |
| glutamate_pos-152.1-59.1_2-4_4/3_normalized fraction | 0.048228385 |
| glutamate_pos-153.1-56.1_2-4_5/0_normalized fraction | 5.42777E-05 |
| glutamate_pos-153.1-57.1_2-4_5/1_normalized fraction | 9.52802E-05 |
| glutamate_pos-153.1-58.1_2-4_5/2_normalized fraction | 0.00089944 |
| glutamate_pos-153.1-59.1_2-4_5/3_normalized fraction | 0.092927696 |

**Raw data and correction**

The chromatogram peaks of 147/75 or 147/41 sometimes have interfering peaks from other metabolites and need to be corrected. The correction is based on the principle that the isotopologue fractions among different set of fragments are identical. For example, the M+1 fraction of glutamate in a sample could be equal to either:

(1) the total ion abundance of 147/74 and 147/75 (negative mode) over the total ion abundance of all 146/74 associated ion pairs
(2) the total ion abundance of 147/102 and 147/103 (negative mode) over the total ion abundance of all 146/102 associated ion pairs,
(3) the total ion abundance of 149/56 and 149/57 (positive mode) over the total ion abundance of all 148/56 associated ion pairs, or
(4) the total ion abundance of 149/84 and 149/85 (positive mode) over the total ion abundance of all 148/84 associated ion pairs.

The corrected fraction of 147/75 is obtained by subtracting the other M+1 fraction (147/74) from the average measured M+1 fractions of other fragmentation associated ion pairs (either the average of bullets 2 and 3, or the average of bullets 2, 3, and 4).

*Corrected fraction of 147/75 = Original fraction of 147/75 - (Fractions of 147/74 and 147/75 - Average of M+1 fractions)*

One example can be found in file Supplementary Data III, sheet S30 in which the 146/74 associated M+1 fractions are compared to 146/102 and 148/84 associated M+1 fractions and the average of the differences is used to correct the ion abundance of 147/75.

**Nonnegative least square regression**

Nonnegative least square regression was implemented using the glmnet R package (Friedman et al., 2010), with the following parameters: lambda = 0, lower.limits = 0, intercept = FALSE and thresh = 1e-30, such that the regression is not regularized, the coefficients are bounded between 0 and 1, and the convergence threshold is small but can generally be reached within the default number of iterations. We compared this method to other R packages with Lawson-Hanson's algorithm (Lawson et al., 1995) implementation such as in the NNLS (Katharine M. Mullen, 2012) or pracma (Borchers, 2021) R packages and found the glmnet-based method to provide the lowest error from our simulation error estimation below. The total fraction of all calculation results should sum to 1. However, regression will produce a value very close to, but not equal to, 1. As a result, normalization is not necessary.

The R script can be found at the GitHub repository (https://github.com/RJDlab/Glu_Asp_Isotopomers).

**Error estimation**

To determine errors in isotopomer distribution estimation due to the use of rank deficient mass isotopomer distribution matrix, we estimated error from the following simulation. We used 5,000 random beta distributions of length 32 for glutamate or 16 for aspartate (eq 1, $\alpha$ set to 2, $\beta$ set to 5) normalized to the sum of 1 to represent positional isotopomer distribution of glutamate or aspartate. For each simulated isotopomer distribution $x_s$, we inferred the MS/MS measurement $T_s$ through the metabolite-specific linear mapping matrix $N$ (eq 2), and then we implemented nonnegative least square to estimate the isotopomer distribution $\hat{x}$ (eq 3). The difference between the simulated isotopomer distribution $x_s$ and the calculated distribution $\hat{x}$ in 5,000 simulations was used to generate the median error and 95% confidence interval (eq 4).

$$x_s \sim Beta\,(\alpha, \beta) \qquad (1)$$

$$T_s = N \cdot x_s \qquad (2)$$

$$\hat{x} = argmin_{x \geq 0} \sum (T_s - N \cdot x)^2 \qquad (3)$$

$$x_{err} = \hat{x} - x_s \qquad (4)$$

The R script can be found at the GitHub repository (https://github.com/RJDlab/Glu_Asp_Isotopomers).

**$F_{C3}$ Calculation using isotopomer distribution**

The $F_{C3}$ calculation is described in the manuscript.

$$F_{C3} = \frac{glu11011}{glu11000 + glu11011}$$

One example can be found in Supplementary Data III, sheet S12, in which the $F_{C3}$ calculated from the isotopomer distribution and the $F_{C3}$ calculated from conventional GLU3, GLU4, and GLU4Q values are compared. The resulting $F_{C3}$ values are similar.

**Estimation of isotopomer distribution errors**

The left charts are the error estimation using matrices with natural abundance correction, and the right one are the error estimation using matrices without natural abundance correction.

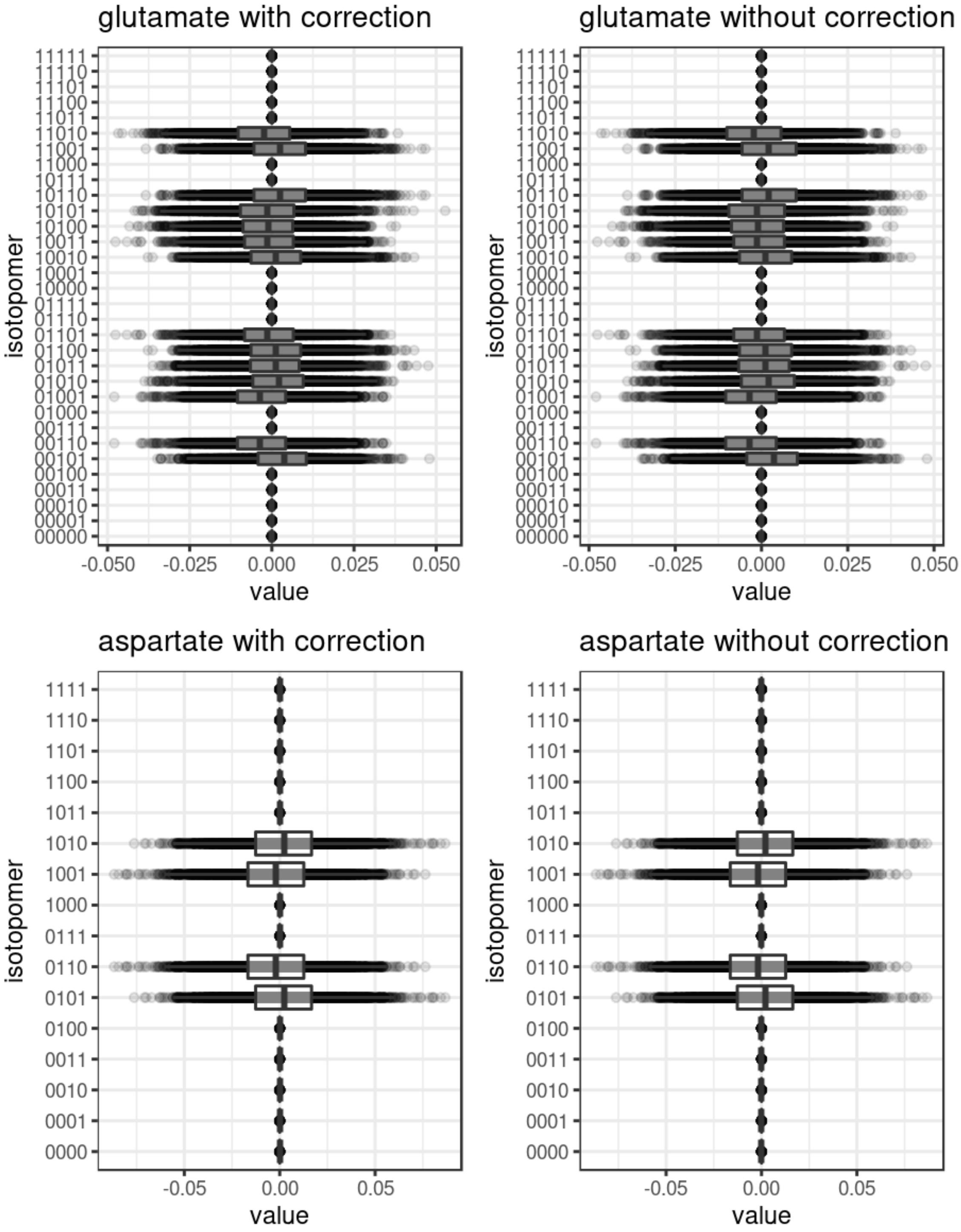
